# Molecular recognition at septin interfaces: the switches hold the key

**DOI:** 10.1101/2020.07.08.161463

**Authors:** Higor Vinícius Dias Rosa, Diego Antonio Leonardo, Gabriel Brognara, José Brandão-Neto, Humberto D’Muniz Pereira, Ana Paula Ulian Araújo, Richard Charles Garratt

## Abstract

The assembly of a septin filament requires that homologous monomers must distinguish between one another in establishing appropriate interfaces with their neighbours. To understand this phenomenon at the molecular level, we present the first four crystal structures of heterodimeric septin complexes. We describe in detail the two distinct types of G-interface present within the octameric particles which must polymerize to form filaments. These are formed between SEPT2 and SEPT6 and between SEPT7 and SEPT3, and their description permits an understanding of the structural basis for the selectivity necessary for correct filament assembly. By replacing SEPT6 by SEPT8 or SEPT11, it is possible to rationalize *Kinoshita’s postulate* which predicts the exchangeability of septins from within a subgroup. Switches I and II, which in classical small GTPases provide a mechanism for nucleotide-dependent conformational change, have been repurposed in septins to play a fundamental role in molecular recognition. Specifically, it is switch I which holds the key to discriminating between the two different G-interfaces. Moreover, residues which are characteristic for a given subgroup play subtle, but pivotal, roles in guaranteeing that the correct interfaces are formed.

**HIGHLIGHTS:**

- High resolution structures of septin heterodimeric complexes reveal new interactions
- Switches of small GTPases are repurposed in septins to play key roles in interface contacts
- The GTP present in catalytically inactive septins participates in molecular recognition
- Conservation of interface residues allows for subunit exchangeability from within septin subgroups
- Specific residues for each septin subgroup provide selectivity for proper filament assembly

**GRAPHICAL ABSTRACT:** 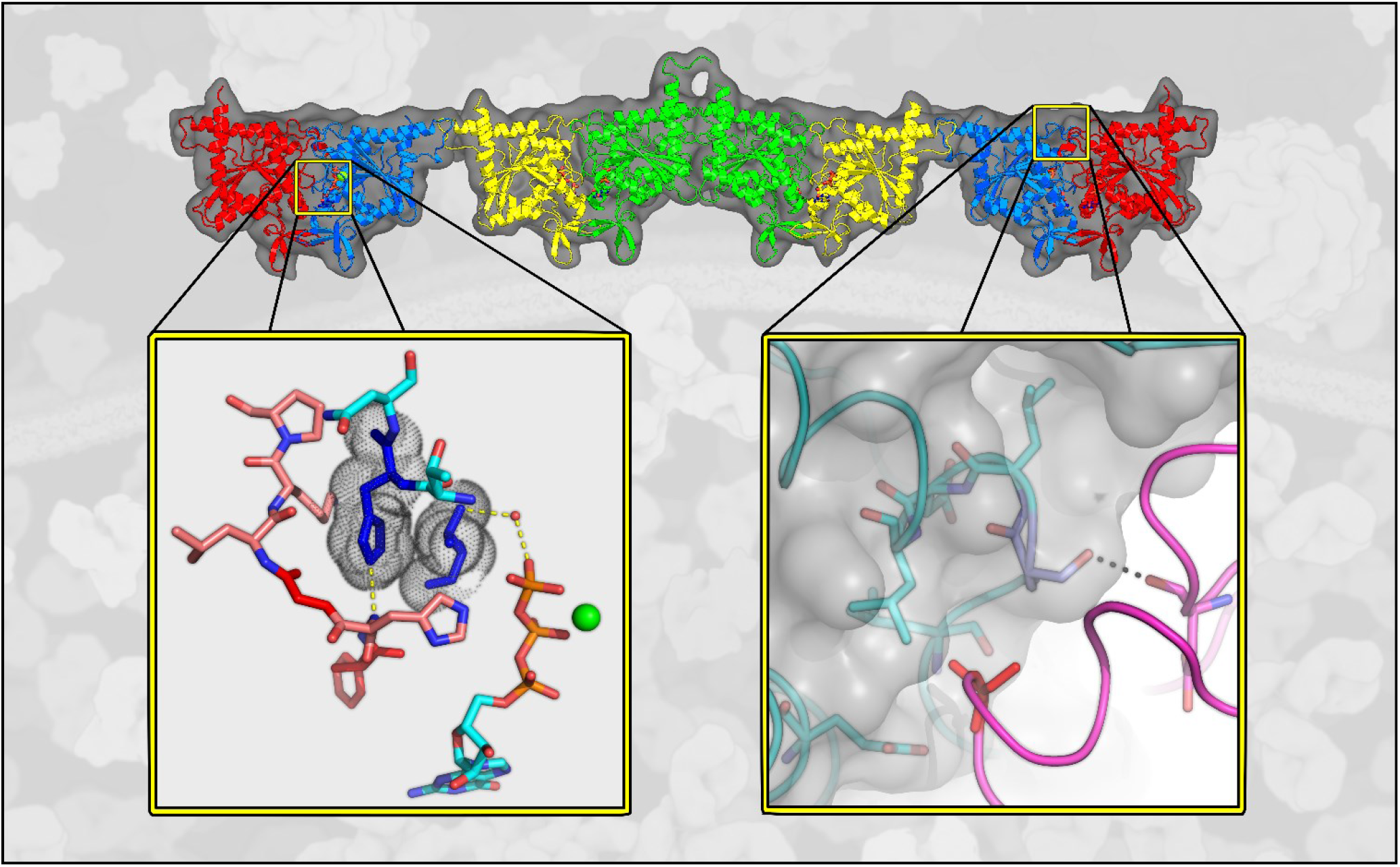

## INTRODUCTION

The assembly of a septin filament is an exquisite but incompletely understood phenomenon. It represents an interesting example of molecular recognition in which different subgroups of homologous molecules must assemble correctly in order to form physiologically functional filaments. These often associate with membranes in order to undertake their roles as molecular scaffolds, membrane modulators, and other cytoskeletal functions[1]. The current view of the septins shows that they are not simply passive molecules acting as scaffolds, but that they act in a physiologically regulated way [2]. Thus, it is not surprising that abnormalities in the expression and function of septins are associated with ageing, infectious diseases, neurodegenerative disorders, male infertility and even cancer [3].

So far, there is no consensus on the existence of septins acting as monomeric units *in vivo*. Rather, their functional form is attributed to filaments and higher-order structures and several septins, when expressed without their partners, are unstable and tend to aggregate [4–6]. Additionally, it has been observed that mutations that prevent the formation of septin filaments in yeast are lethal, showing the importance of their organization into complex structures [7]. A common characteristic observed in all such complexes is the linear arrangement of monomers into filaments to form non-polar polymers [8–11]. These present two-fold and pseudo two-fold symmetry axes perpendicular to the main filament axis rather than a screw axis parallel to it.

Thirteen human septins can be divided into four subgroups based on their domain structure and sequence similarity. These are named the SEPT2 (SEPT1, 2, 4, and 5), SEPT3 (SEPT3, 9 and 12), SEPT6 (SEPT6, 8, 10, 11, and 14), and SEPT7 subgroups, and they assemble into filaments based on the polymerization of core particles. The latter can be either dimers of tetramers (involving the participation of one septin from each subgroup) or dimers of trimers (in which the SEPT3 subgroup is missing). It has recently been established that the correct order for the subgroup septins in the resulting octamers and hexamers is SEPT2-SEPT6-SEPT7-SEPT3-SEPT3-SEPT7-SEPT6-SEPT2 and SEPT2-SEPT6-SEPT7-SEPT7-SEPT6-SEPT2 respectively [12,13] and it is believed that subgroup members should be exchangeable. Here we refer to this important prediction as *Kinoshita’s postulate*, after the original proposal made in 2003 [14]. If this postulate is strictly applied it could give rise to 60 different types of octamer and 20 types of hexamer. Several, but far from all of these possibilities, have been reported to assemble either *in vivo* or *in* [15–18].

The correct assembly of such linear filaments requires that each septin monomer provides two different interfaces through which it interacts with its immediate neighbours. These are labelled G- and NC- because they employ the guanine nucleotide-binding site (G-interface) or the N- and C-terminal extensions (NC-interface). Although these auxiliary N- and C-terminal domains are believed to play a role in assembly [10,19], it is contacts made between the G-domain which dominate our current view of the filament, particularly at the G-interfaces.

Based on the above, it is expected that a filament based purely on octamers would present the following five interface types: SEPT3-SEPT3 (NC), SEPT3-SEPT7 (G), SEPT7-SEPT6 (NC), SEPT6-SEPT2 (G) and SEPT2-SEPT2 (NC). The latter arises when two octamers associate by end-to-end interactions during polymerization. Hexamers, on the other hand, would lack both the SEPT7-SEPT3 and SEPT3-SEPT3 interfaces but acquire a new one in their place (a SEPT7-SEPT7 G-interface).

In order to fully understand the molecular basis of spontaneous filament assembly, each of the six different interfaces needs to be characterized at the atomic level. In this context, it is curious that when the G-domains of individual septins are crystallized, the vast majority form filaments within the crystal which employ the same G- and NC-interfaces, even if these are not anticipated from the canonical organization of the oligomers. This can be rationalized by appreciating the considerable sequence similarity which exists between the G-domains of different subgroups allowing for the formation of promiscuous interfaces. However, this raises an important question; how is each monomer able to identify its correct position within the oligomer? Individual septins have been extensively characterized showing their ability to bind guanine nucleotides [10,20–22]. However, few studies have focused on the interface details of the mammalian heterocomplexes [4,5,19,22]. To contribute to a fuller understanding of both biochemical and structural aspects that govern the interaction selectivity amongst septin subgroups (*Kinoshita’s postulate*), here we have co-expressed and characterized 12 septin heterodimeric G-domain complexes including representatives of all four human septin subgroups.

Attempts to crystallize heteromeric septin complexes have met with little success. Only the X-ray structure of the SEPT2-SEPT6-SEPT7 complex has been described to date and, even that is substantially incomplete due to its limited resolution [10]. Although a landmark in providing insight into the structure of the filament itself, it provides very little information about the atomic details of the interfaces and the molecular determinants of selectivity. In order to overcome these difficulties we have taken a “divide and conquer” approach in which we aim to structurally interrogate each interface individually [20,23]. Here we apply this approach to the 12 heterodimeric complexes and report the crystal structures of four. In so doing, we report on details of the interactions which stabilize the G-interfaces, provide a molecular basis for Kinoshita’s postulate and highlight the unexpected but pivotal role played by switches I and II in controlling inter-subunit selectivity.

## RESULTS AND DISCUSSION

### Differential stability among complexes

Details of the heterologous expression and purification of all 12 heterodimeric complexes, along with the homodimer of SEPT7, are given in Table S1. The SEC-MALS profiles of all complexes show a predominant monodisperse peak, consistent with the molecular mass of a dimer (Fig. S1, Table S2). This oligomeric state was expected as all chosen pairs are anticipated to form G-interface dimers and the absence of the N- and C-terminal domains disfavours higher-order states. The sole exceptions were the constructs of the SEPT3 subgroup (SEPT3 and SEPT9), which include the C-terminal domain. However, these are rather small and do not form coiled-coils like the remaining subgroups. Nucleotide-content assays confirmed the presence of natively-bound nucleotides for all of the purified heterodimers. All samples contained GDP whilst only those containing a catalytically-inactive SEPT6 subgroup-member (SEPT6, SEPT8 and SEPT11) also contained GTP, in roughly equal amounts (Fig. S2). Based on these findings and on previous reports [10,22,24,25], it is readily inferred that these dimers form by employing nucleotide-loaded G-interfaces.

An analysis of the thermal-stabilities of the different heterodimeric complexes (Fig. 1) shows that, regardless of the intrinsic stabilities inherent to each monomer, combinations involving members of the SEPT2 and SEPT6 subgroups (shown in blue, red and yellow) are more stable than the two heterodimers involving SEPT7 (SEPT3-SEPT7 and SEPT9-SEPT7, shown in pale green). However, a SEPT3 mutant (SEPT3_T282Y_), which recovers a G-interface tyrosine present in all other subgroups, forms a more stable dimer with SEPT7 than does the wild-type. Indeed, the observed inflection temperature (Ti) is similar to that of the complexes involving SEPT4 and SEPT5. Similarly, SEPT7G alone also forms a slightly more stable homodimer than the SEPT7-SEPT3 complex (Fig. 1, dark green). It is interesting to note that the three complexes formed by SEPT2 (with SEPT6, SEPT8 or SEPT11) are more thermostable than similar complexes formed by the remaining SEPT2 subgroup members (SEPT4 or SEPT5) and in all cases that formed with SEPT8 is the least stable of all. To understand these observations more fully, we embarked upon a systematic attempt to crystallize the heterodimers and determine their structures.

**Fig. 1.**
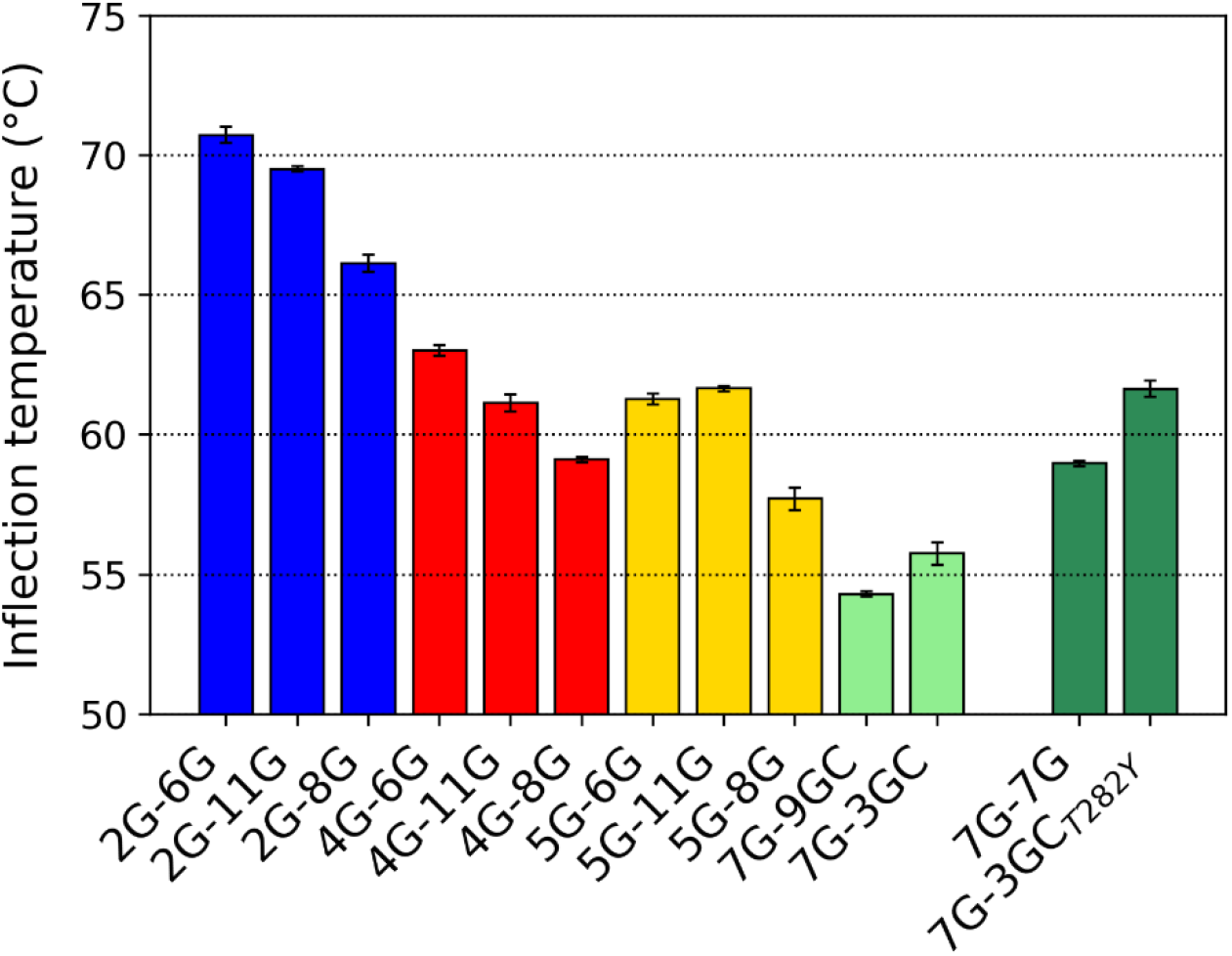
Thermal stability using intrinsic fluorescence (Tycho). The Inflection temperature (Ti) is given for all 12 heterodimers as well as the SEPT7G homodimer. Error bars are derived from three independent measurements. Heterocomplexes involving members from SEPT2 and SEPT6 subgroups are more stable than those involving members from SEPT7 and SEPT3 subgroups.

### Overall description of the dimers

Of the 12 heterodimeric complexes which were successfully expressed and purified, four produced X-ray diffraction quality crystals. These include at least one representative member from each of the four different subgroups. Specifically, the complexes described here are SEPT2-SEPT6, SEPT2-SEPT8, SEPT2-SEPT11 and SEPT7-SEPT3_T282Y_, in which SEPT6, SEPT8 and SEPT11 all belong to the SEPT6 subgroup and will be referred to collectively by the shorthand SEPT6/8/11. Henceforth, the specification of the domain structure of the construct, G or GC, will be dropped. Table 1 gives a summary of the crystallographic data collection and refinement statistics for all of the structures. The resolutions vary from 1.86 Å for SEPT2-SEPT11 to 2.75 Å for SEPT7-SEPT3_T282Y_.

**Table 1.** Crystallographic data collection and refinement statistics. Values in parentheses refer to the highest resolution shell.

|  | SEPT2-SEPT11 | SEPT2-SEPT6 | SEPT7-SEPT3<sub>T282Y</sub> | SEPT2-SEPT8 |
| --- | --- | --- | --- | --- |
| *PDB ID* | **6UPQ** | **6UPA** | **6UQQ** | **6UPR** |
| X-ray Source | Diamond I24 | Diamond I24 | Diamond I24 | Diamond I04 |
| Detector | Pilatus3 6M | Pilatus3 6M | Pilatus3 6M | Eiger2 X 16M |
| Cell parameters (Å) a; b; c;<br>$\alpha$; $\beta$; $\gamma$ | 56.66, 81.86, 68.22<br>90.00, 97.14, 90.00 | 82.73, 82.73, 170.35<br>90.00, 90.00, 90.00 | 61.39, 85.53, 152.05<br>90.00, 90.97, 90.00 | 52.74, 82.80, 78.17<br>90.00, 103.63, 90.00 |
| Space Group | $P2_1$ | $P4_12_12$ | $P2_1$ | $P2_1$ |
| Resolution (Å) | 86.86 – 1.86 (1.91 – 1.86) | 37.21 – 2.51 (2.58 - 2.51) | 61.38 – 2.75 (2.87 – 2.75) | 82.80 – 2.30 (2.36 – 2.30) |
| $\lambda$ (Å) | 0.96863 | 0.96861 | 0.96863 | 0.97950 |
| Multiplicity | 5.7 (6.8) | 12.8 (13.3) | 3.6 (3.6) | 6.9 (7.1) |
| R<sub>pim</sub> (%) | 8 (129.2) | 4.4 (63.5) | 9.8 (54.8) | 5.5 (68.0) |
| CC(1/2) | 0.973 (0.453) | 0.999 (0.564) | 0.988 (0.604) | 0.992 (0.782) |
| Completeness (%) | 99.6 (98.9) | 99.9 (99.9) | 94.8 (96.4) | 99.2 (99.5) |
| Reflections | 347126 (25676) | 269502 (20143) | 139027 (17068) | 119518 (14951) |
| Unique Reflections | 51728 (3798) | 21021 (1516) | 38827 (4789) | 28853 (2108) |
| $I/\sigma$ | 5.8 (0.9) | 11.6 (1.2) | 5.7 (1.3) | 8.0 (1.0) |
| Reflections used for<br>Refinement | 51621 | 20952 | 38766 | 28678 |
| R (%)** | 18.8 | 22.41 | 19.70 | 20.88 |
| R<sub>free</sub> (%)** | 21.8 | 26.55 | 25.55 | 25.55 |
| N° of protein atoms | 4148 | 3962 | 8191 | 4317 |
| N° of ligand atoms | 61 | 61 | 112 | 61 |
| B (Å<sup>2</sup>) | 37.03 | 33.05 | 61.61 | 32.93 |
| Coordinate Error (ML based)<br>(Å) | 0.27 | 0.41 | 0.44 | 0.39 |
| Phase error (°) | 24.59 | 27.24 | 28.52 | 33.88 |
| *Ramachandran Plot* |  |  |  |  |
| Favoured (%) | 96.87 | 96.15 | 94.86 | 95.10 |
| Allowed (%) | 2.74 | 3.62 | 5.14 | 4.71 |
| Outliers (%) | 0.39 | 0.20 | 0.00 | 0.19 |
| All-atom Clashscore | 2.87 | 2.77 | 4.81 | 4.51 |
| *RMSD from ideal geometry* |  |  |  |  |
| r.m.s. bond lengths (Å) | 0.003 | 0.002 | 0.003 | 0.003 |
| r.m.s. bond angles (°) | 0.687 | 0.502 | 0.559 | 0.591 |

The septin fold has been described previously [6,10] and is based on the architecture of an αβα three-layered sandwich [26]. The fold is dominated by a central six-stranded β-sheet surrounded by α-helices, followed by the *septin unique element.* None of the structures described here deviate significantly from the canonical septin fold and we will adopt the standard nomenclature used for the elements of secondary structure (Fig. S3).

Given the recently established order for septins within the octameric core particle [12,13], all of the four complexes reported here are expected to present physiological heterodimeric G-interfaces. This is based on Kinoshita’s proposal that members of a given subgroup should be interchangeable such that SEPT6, SEPT8 and SEPT11 should form an equivalent interface with SEPT2. In the case of the complex formed between SEPT7 and SEPT3, it was necessary to employ the T282Y mutant of the latter which has been shown to stabilize G-interface homodimers of SEPT3 [25]. All crystallization attempts of complexes containing native SEPT3 yielded crystals of SEPT7 alone.

The three complexes formed with SEPT2 have G-interface dimers in the asymmetric unit. Several structures are available for SEPT2 including both in isolation (PDB IDs 2QNR, 2QA5, 3FTQ) and as part of the SEPT2-SEPT6-SEPT7 heterotrimeric complex (PDB ID 2QAG) [10,27]. The latter also includes the only currently available structure for SEPT6. Fig. S4 shows that the structures provided here for both subunits are significantly more complete than those described previously. Most notable is that switch I is fully ordered in SEPT2 and is almost complete in SEPT6, where only two residues present poor electron density. This is in direct contrast to that observed for the SEPT2 structures in isolation where this region systematically presents considerable disorder. This seems to imply that the correct physiological pairing of septins across the G-interface has a profound effect on the conformation of the switch.

The remaining two complexes containing SEPT2 provide the first structural information for both SEPT11 and SEPT8. In both cases, the switch I region of each component of the dimer shows an effectively identical conformation to that seen with SEPT2-SEPT6. However, the conformations observed for each subunit of the complex are significantly different, including the presence of a short α-helix in the case of SEPT2 and a five-residue deletion in all of the SEPT6 subgroup members (Fig. S4c). Indeed, the overall structures of the three dimers are all extremely similar, with the relative orientation of the two subunits being almost indistinguishable and yielding a mean RMSD of 0.29 Å on C^α^ atoms (Fig. 2a). This suggests not only a favourable complementarity between the two subunits but also that any SEPT6 subgroup member is accommodated well by the interaction surface provided by SEPT2, in accordance with Kinoshita’s postulate [14].

**Fig. 2.**
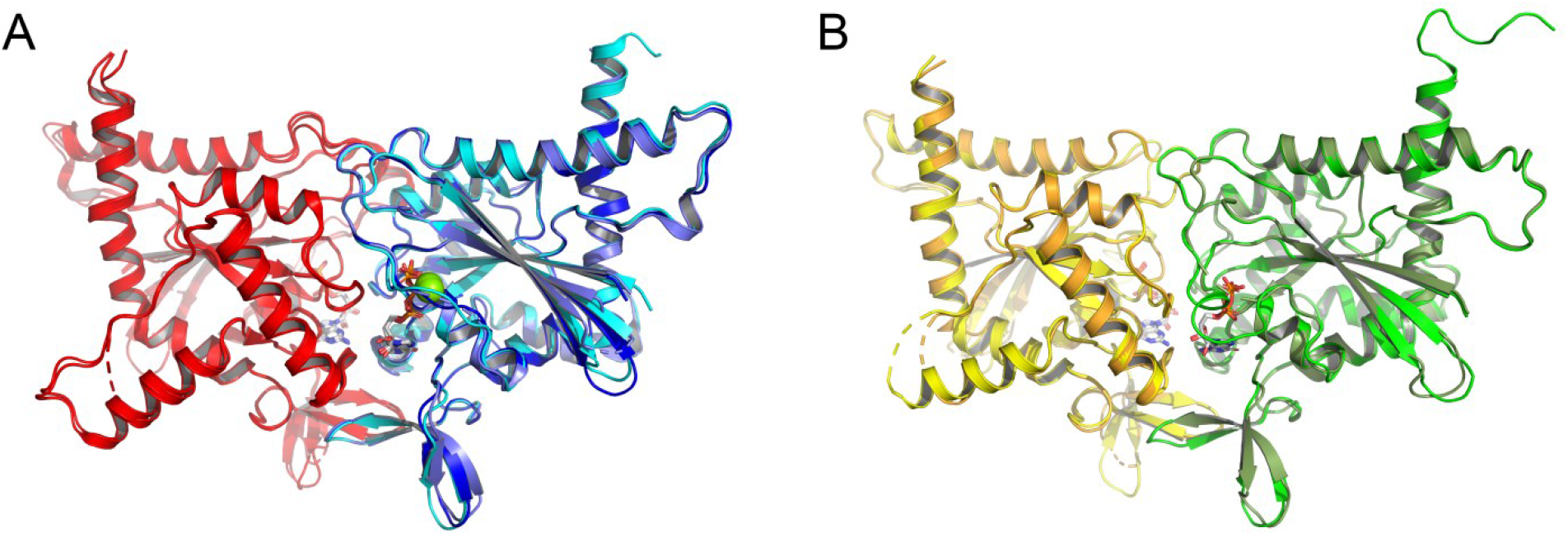
Global superposition of G-interface dimers. (a) dimers formed between SEPT2 and members of the SEPT6 subgroup. In all cases, SEPT2 is shown in red. SEPT11 (dark blue), SEPT8 (light blue) and SEPT6 (cyan) are almost indistinguishable. (b) the two crystallographically independent G-interface dimers formed between SEPT7 (shades of yellow) and SEPT3_T282Y_ (shades of green). Variations are minimal. The colour scheme used to represent the four different subgroups will be preserved, as much as is possible, in the subsequent figures.

In the case of the SEPT7-SEPT3_T282Y_ complex, there are four monomers in the asymmetric unit. These form a SEPT7 NC-homodimer (chains A and B) and a SEPT3_T282Y_ NC-homodimer (chains C and D). Crystallographic symmetry can be used to generate two independent G-interface dimers which superpose extremely well, yielding an RMSD of 0.33 Å (Fig. 2b). Slightly higher values (0.74 Å on average) are obtained on comparison with the three SEPT2 containing complexes. For both of the individual components (SEPT7 and SEPT3), high resolution structures are already available [20,23]. The structure of SEPT7 alone (PDB ID 6N0B) superposes extremely well with the SEPT7 subunit of the heterodimer, yielding a mean RMSD of 0.36 Å for eight crystallographically independent comparisons. SEPT3 superposes less well on its counterpart in the complex (mean RMSD of 1.1 Å) principally due to differences in the switch regions. In this complex, there is no systematic gain in the structure of switch I as observed for the three complexes made with SEPT2. Both SEPT7 and SEPT3 present some subunits with a complete switch I and others present some degree of disorder. However, a notable difference between them is that the SEPT7 subunits which present a complete switch I have identical conformations independent of being part of the complex or not (Fig. S5). This is not the case for SEPT3_T282Y_ where there is much greater variation.

The nucleotides observed in each of the subunits of each complex (GTP in the case of SEPT6, SEPT8 and SEPT11 and GDP in the case of SEPT2, SEPT7 and SEPT3_T282Y_) are coherent with the nucleotide content described above (Fig. S2) and with the lack of catalytic activity observed previously for the SEPT6 subgroup [10,22,28]. There is little of note about the nucleotide-binding sites and Mg^2+^ coordination when compared to that described previously [23,27–29], with the exception of Arg66 from switch I in SEPT2 which interacts with the α and β phosphates of the GDP. This arginine, which is disordered in previous crystal structures, is *characteristic* of the SEPT2 subgroup (see below) and interacts with the nucleotide in an identical fashion in all three complexes. This is presumably related to the complete ordering of the entire switch due to specific interactions made with SEPT6/8/11, as described below.

### Filaments are observed in all cases

By generating symmetry-related molecules it is possible to observe the presence of filaments in all of the crystal structures reported here. These present alternating NC- and G-interfaces typical of the majority of septin structures reported to date. Here, we focus on the G-interfaces which are formed between SEPT2 and SEPT6/8/11 and between SEPT7 and SEPT3_T282Y_. These are all physiological (Fig. 3a) and are readily observed in all of the crystal structures (Fig. 3b and c). Based on the model given in Fig. 3a it might be anticipated that two monomers of SEPT2 would pack to form a physiological NC-interface. However, this is not observed in any of the structures involving SEPT2. Rather, an NC-interface is made with a neighbouring copy of SEPT6/8/11 and the two different monomers, therefore, intercalate along the filament. Since these interfaces are not expected to arise in a physiological octamer or hexamer-based filament, they are considered to be promiscuous. The largest physiological entity formed in this case is, therefore, the G-interface heterodimer (Fig. 3b). The lack of a homotypic NC-interface between two SEPT2 monomers is likely the result of the construct used in this work which lacks the N-terminal domain. This is known to embrace its partner across the NC-interface in a “domain-swapped” arrangement [30]. Its absence may destabilize the interface to the point of disfavouring it over the promiscuous SEPT2-SEPT6 interaction. Although it is difficult to see from the structure why the latter would be favoured, this observation seems to be consistent with recent work showing that the SEPT2-SEPT2 NC-interface is the most fragile along the filament (at least with respect to salt concentration) and that is why SEPT2 is observed at the end of the core particles (Fig. 3a). It is also consistent with work in yeast using particles incorporating a mutant of septin cdc11 lacking the N-terminal domain. Although this resulted in octamers, with the mutant protein occupying the terminal position, these were unable to polymerize into filaments [11].

**Fig. 3.**
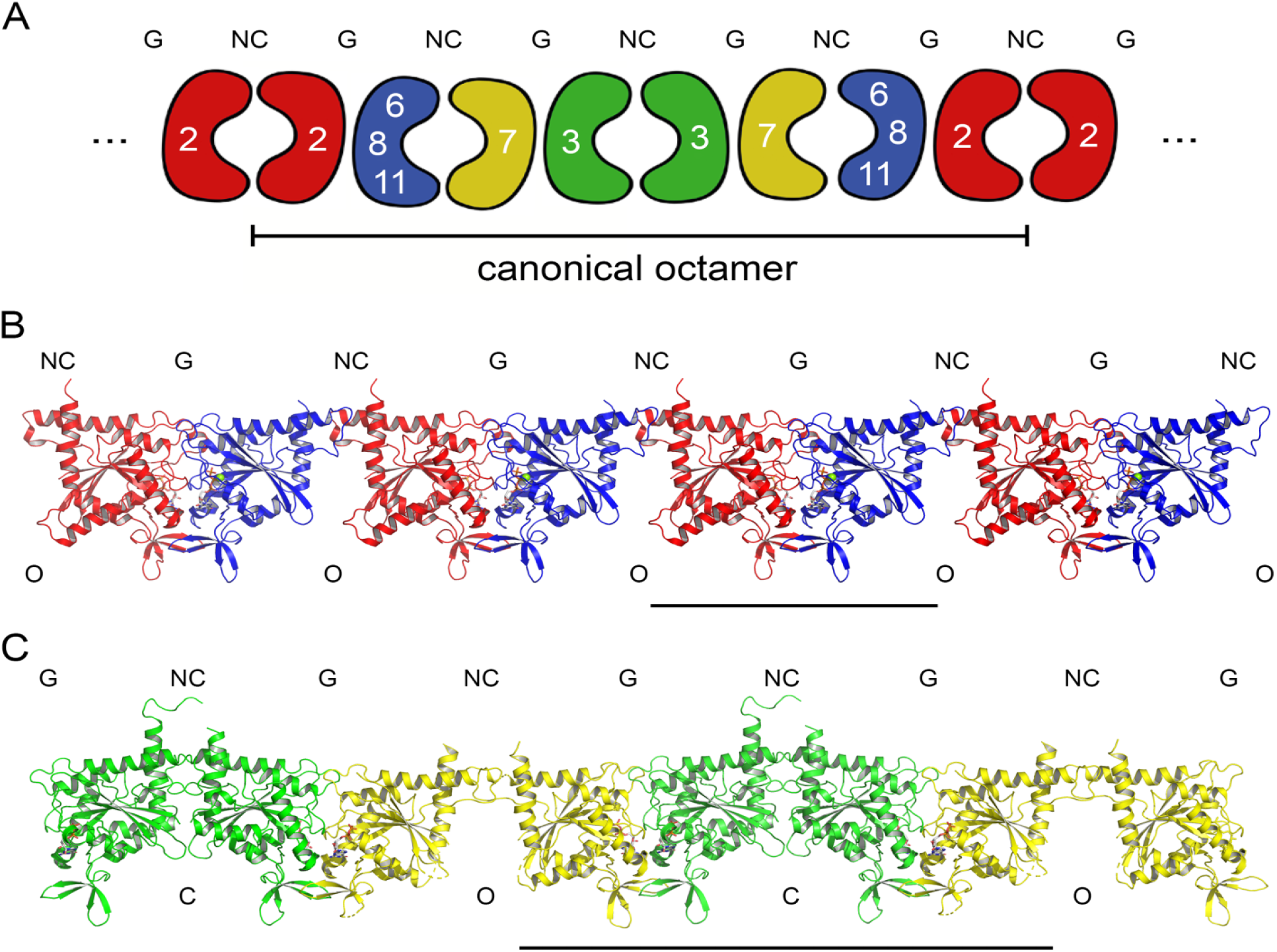
Filaments form in all crystal structures. (a) shows the canonical structure for a filament based on octamers which terminate with SEPT2. End-to-end contacts between two copies of SEPT2 at an NC-interface allows the octamers to polymerize. (b) A filament formed by the SEPT2-SEPT11 complex in red and blue respectively. SEPT2-SEPT8 and SEPT2-SEPT6 are essentially identical. The solid bar indicates the G-dimer which is present as part of the canonical arrangement shown in (a). (c) A filament formed by the SEPT7-SEPT3_T282Y_ complex. The solid bar indicates a tetrameric unit which includes a sequence of three interfaces anticipated by the canonical model shown in (a). These tetramers are united by a non-canonical NC-interface formed between two copies of SEPT7.

The SEPT7-SEPT3_T282Y_ filament is rather different, with the two septins appearing as alternating dimers (Fig. 3c). Besides the G-interface formed between SEPT7 and SEPT3_T282Y_, a second physiological interface (a homotypic NC-interface between two SEPT3_T282Y_ monomers) arises as a result of crystallographic symmetry (Fig. 3a). As a consequence, a sequence of four successive monomers is present exactly as anticipated in the octameric particle (Fig. 3a). These tetramers include three physiological interfaces (one homotypic NC-interface and two heterotypic G-interfaces) which are united by promiscuous NC-interfaces formed between two monomers of SEPT7. The homotypic NC-interfaces between SEPT7 monomers are in the canonical open (O) conformation whilst those between SEPT3_T282Y_ monomers are squeezed together in the closed (C) conformation [23]. The latter is discussed more fully below.

### The G-interfaces reveal a critical role for switch I

Since the description which follows will focus mostly on the nature of the interfaces themselves and the origin of selectivity in filament assembly, it is of use to identify residues which are characteristic of each of the four subgroups. These are defined as residues present in all members of a given subgroup of human septins but absent from all others (Fig. S3). Fig. S6 shows that several of these *characteristic* residues in the SEPT2 and SEPT6 subgroups map to the G-interface between them. To produce representative figures, the structure of the complex formed between SEPT2 and SEPT11 will generally be used, as it is of higher resolution, but the equivalent sequence numbers for SEPT6 will also be quoted, as it is considered the gold-standard for the subgroup.

In Fig. 4, the G-interfaces for each of the complexes formed by SEPT2 have been opened like a book and the subunits separated in order to readily visualize the regions of the structure which make direct contact across the interface. There is a remarkable similarity in the patterns generated when mapping contact residues onto the molecular surface in all three cases. This illustrates clearly why different members of the SEPT6 subgroup are able to substitute for one another in pairing with SEPT2 and provides a structural basis for Kinoshita’s postulate [14]. There is almost complete conservation of the residues involved in forming the core of the interface with the exception of a small number of conservative substitutions. However, some differences do arise in the periphery of the contact area. These are often highly exposed residues which present side chain conformational variation, some of which is probably due to variation in the resolution of the three structures. Overall, they appear to be of little significance, with the only possible exception being a new salt-bridge in the SEPT8 complex which arises as a consequence of the substitution of Pro70 in SEPT11 (Pro71 in SEPT6) by Glu73 in SEPT8. This leads to an inter-chain hydrogen bond with Arg198 of SEPT2. Finally, the presence of His77 only in SEPT8 is notable but provides no strong new interaction.

**Fig. 4.**
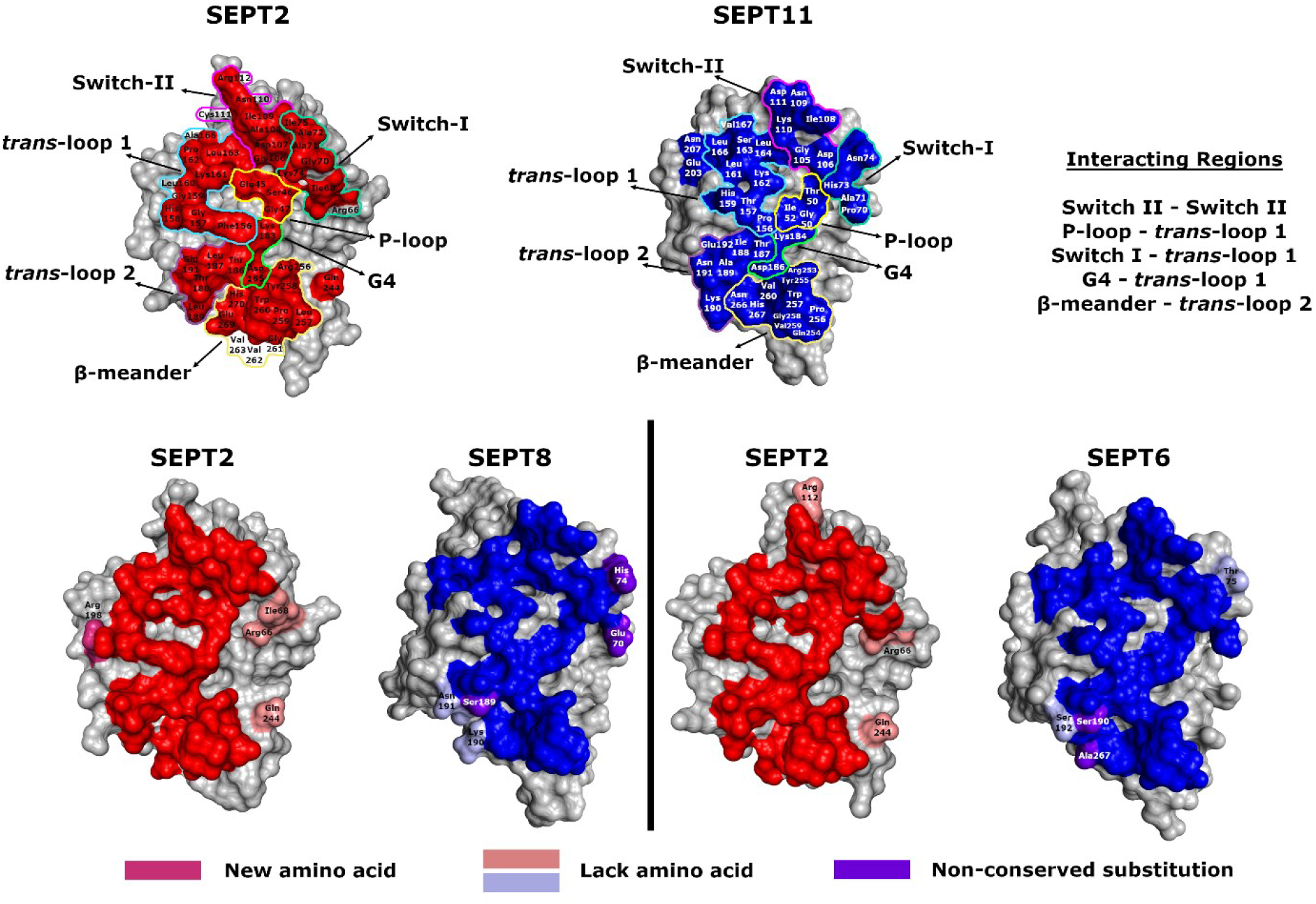
The interfaces for the SEPT2-SEPT6/8/11 complexes. The subunits have been separated and rotated by 90° around the y-axis to expose the interface residues (red and blue respectively for SEPT2 and SEPT11, taken to be the reference structure.) Additional colours follow the index at the bottom of the figure and apply to the SEPT8 and SEPT6 complexes when compared to that of SEPT11. Pairs of contact regions are indicated (top right). The interactions at the interface’s core of these complexes are maintained, despite some variance in rim residues.

Seven regions of the surface are involved in the core contacts. These are switch I, switch II, the P-loop, G4, the *trans* loops 1 and 2 [31], and the β-meander. They map onto one another across the interface in accordance with the information given in Fig. 4. The only region which maps onto itself is switch II where an anti-parallel wide-type β-bridge forms symmetrical hydrogen bonds across the interface [20,24,32]. Although classically associated with establishing the preference for the guanine base, the G4 motif in septins also forms part of the G-interface, where Lys183 in SEPT2 reaches across the interface to form a hydrogen bond with the carbonyl from a conserved proline of the *trans* loop 1. However, the most noticeable region of contact is that made by switch I of each subunit with the corresponding *trans* loop 1 (which connects the β4 strand to α3). As mentioned above the switch I region is often largely disordered, particularly in GDP-bearing structures. However, it is not only completely ordered in all subunits of the SEPT2 complexes (with the exception of two residues in SEPT6) but forms a significant part of the buried surface area and generates selective interactions which probably explain why the SEPT2 and SEPT6 subgroups select one another in forming a native G-interface (see below).

On average, switch I from SEPT2 is responsible for forming 10.6% of the contact surface with an additional 7.2% coming from the switch I region of SEPT6/8/11. Together, the two switches are responsible for burying over 350 Å^2^ of surface area on dimer formation, forming a significant portion of the interface (17.8%). This is in stark contrast to the SEPT7-SEPT3_T282Y_ dimer where, besides having an overall smaller contact area (1,717 Å^2^ compared with 2,033 Å^2^), the sum contribution of switch I from both subunits is only 3%. This is evident in Fig. 5 where it can be seen that the switch I region contributes only marginally to the interface in SEPT7 (Ser84/His85) and not at all in SEPT3_T282Y_, despite being fully ordered in both cases.

**Fig. 5.**
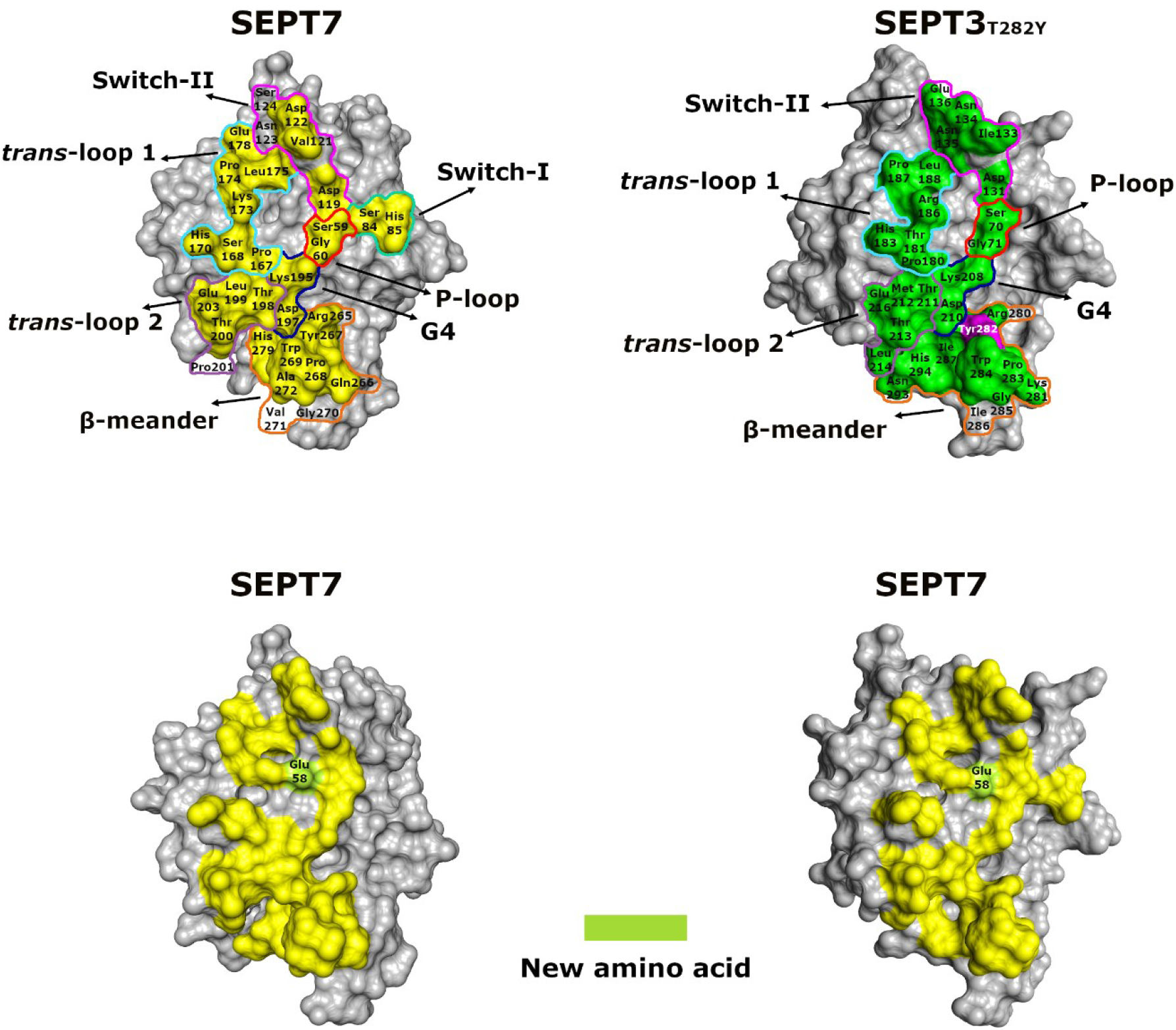
The dimeric interfaces made by SEPT7. Yellow and green respectively indicate contact residues in the SEPT7-SEPT3_T282Y_ complex. The mutated residue (T282Y) is shown in purple. For comparison, contact residues involved in the homodimeric interface formed by two copies of SEPT7 (below) are shown in yellow (from PDB ID 6N0B). In both cases, switch I barely participates in the interface.

This notable difference between the two types of native G-interface (SEPT2-SEPT6/8/11 on the one hand and SEPT7-SEPT3_T282Y_ on the other) becomes clearer on superposing the two types of dimer (Fig. 6). Whereas switch I of SEPT2 and SEPT6/8/11 are shifted towards the interface and contribute significantly to direct contacts, this is not the case with SEPT7 and SEPT3_T282Y_. This additional buried surface area is expected to be a major contributor to the greater thermal stability of the SEPT2 heterodimers as mentioned above (Fig. 1). The SEPT2 complexes present T_i_ values between 66.1 and 70.7 °C, all significantly higher than the 55.7 °C observed for SEPT7-SEPT3. The reduced stability of the latter is probably a characteristic of this particular subgroup combination, since SEPT7-SEPT9 has a similar T_i_ value of 54.3 °C.

**Fig. 6.**
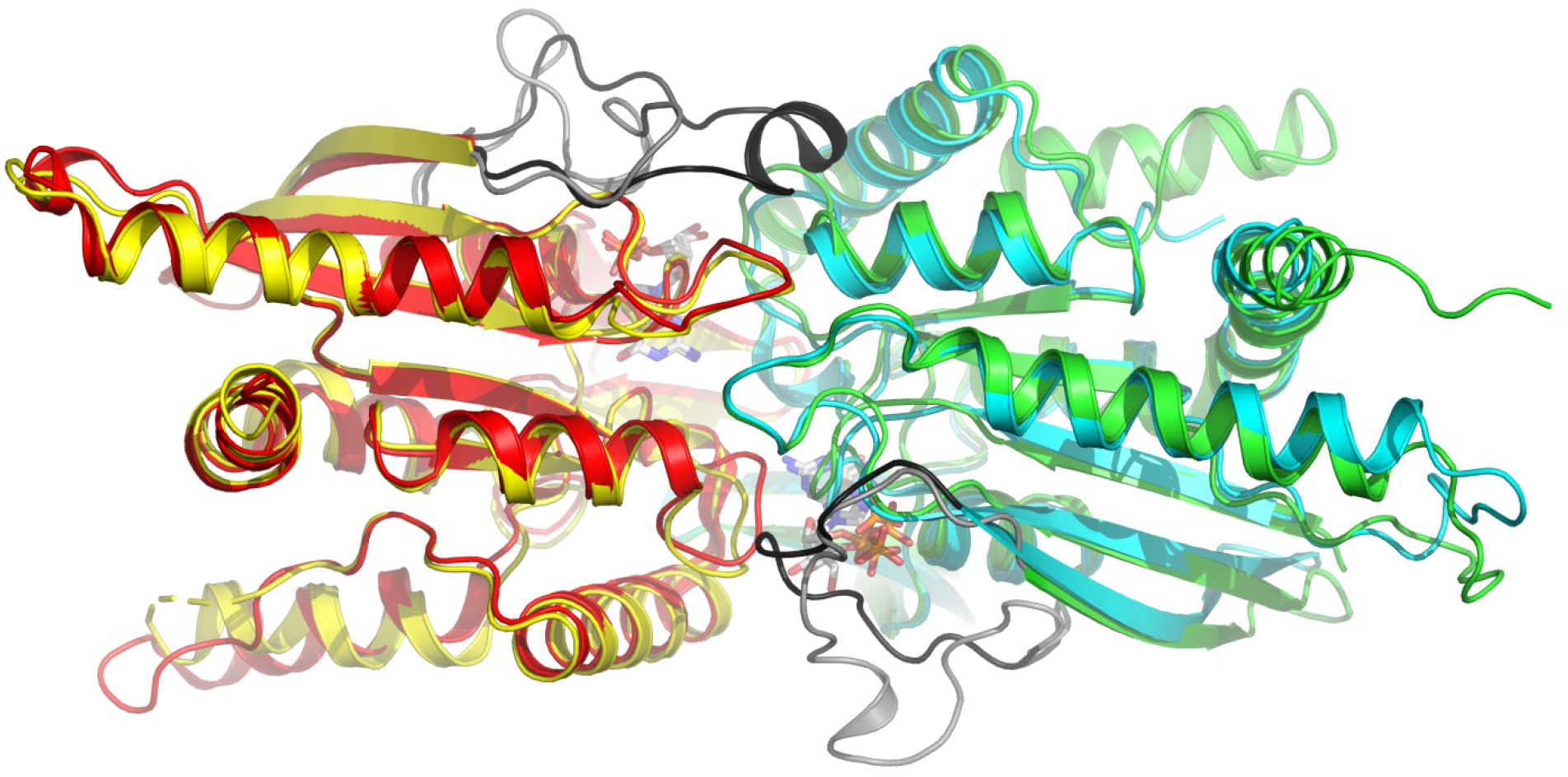
Superposition of G-interface heterodimers. The SEPT2-SEPT11 dimer is shown in red/blue and that of SEPT7/SEPT3_T282Y_ in yellow/green. The corresponding switch I regions are shown in black and grey respectively. Those from both SEPT2 and SEPT11 approach the partner subunit much more closely than observed for SEPT7 and SEPT3T282Y.

A similar situation is observed in the homodimeric structure of SEPT7 (PDB ID 6N0B) where the two switch I regions are responsible for only 2.9% of the contact area. Indeed, the contact surfaces of both the homodimeric and heterodimeric structures (Fig. 5) are very similar which is consistent with the formation of both such G-interfaces in physiological oligomers; SEPT7-SEPT7 in the case of hexamers and SEPT7-SEPT3 in the case of octamers. Consistent with this, the SEPT7 homodimer has a similar thermal stability (T_i_ = 58.9 °C) to that of the heterodimer with SEPT3. The apparently reduced stability of these complexes may be related to the need to form both types of oligomer *in vivo*, for which an overly stable interface may be disadvantageous. This would permit the formation of mixed filaments formed of both oligomeric particles [13].

Several features could contribute to the fine-tuning of the subunit affinities between SEPT7 and its possible partners: either SEPT3 or with a second copy of SEPT7. Most notable is a symmetrical array of salt-bridges across the interface in the homodimer (involving Glu58 and Lys173 from both subunits) which is compromised in the heterodimer due to the substitution of Glu58 by the *characteristic* residue Gln69 in SEPT3. This eliminates one negative charge from the array and leads to structural disorder. This may explain the slightly greater thermal stability of the SEPT7 homodimer. Nevertheless, it would seem that both possibilities are viable but neither shows the thermal stability found for the heterocomplexes involving SEPT2.

The mutation in SEPT3 introduced a tyrosine into the β-meander of the G-interface. This led to an increase of about 6 °C in the thermal stability of the heterodimer (Fig. 1) and the tyrosine forms an equivalent interaction across the interface to that observed in all three SEPT2 complexes as well as the homodimer of SEPT7 (Fig. S7). The hydrogen bond formed with Asp197 of SEPT7 probably explains the increase in thermal stability, at least in part. It is interesting to note that when coexpressing the equivalent yeast septins, cdc3 and cdc10, these fail to interact unless in the presence of cdc12 [33] implying that there may be a fragile interface at an equivalent position along the oligomer in both systems.

### Switch I of SEPT6/8/11 plays a pivotal role

The reason that switch I of SEPT6/8/11 is ordered in the three heterodimers is because specific interactions are made with SEPT2. In order to build selectivity into the interface, it might be expected that this would involve the participation of *characteristic* residues coming from the two different subunits. His73 (His74 in SEPT6) from switch I is such a residue and interacts with *trans* loop 1 of SEPT2. As can be seen in Fig. 7a, *trans* loop 1 adopts a unique conformation in SEPT2, when compared to the remaining subgroups, and the main chain from Gly157 to Lys161 provides a cavity into which the side chain of His73 can nestle snuggly (Fig. 7b). Gly159, which forms part of the cavity, is not strictly a *characteristic* residue for the SEPT2 subgroup but is almost so (Fig. S3) and has a highly extended conformation (φ = 178, Ψ=-160 in SEPT2-SEPT11). The resulting difference to the conformation of *trans* loop 1 appears to be the result of Gly159 together with Phe156 which is a *characteristic* residue for the SEPT2 subgroup and reaches down into the centre of the interface. The only specific interaction made by the side chain of His73 of switch I is by accepting a H-bond via N_ε2_ from the main chain amide of His158 (Fig. 7b) suggesting the histidine to be in the less common π tautomeric form. This interaction would be impossible with the loop conformation taken by any of the other subgroups.

**Fig. 7.**
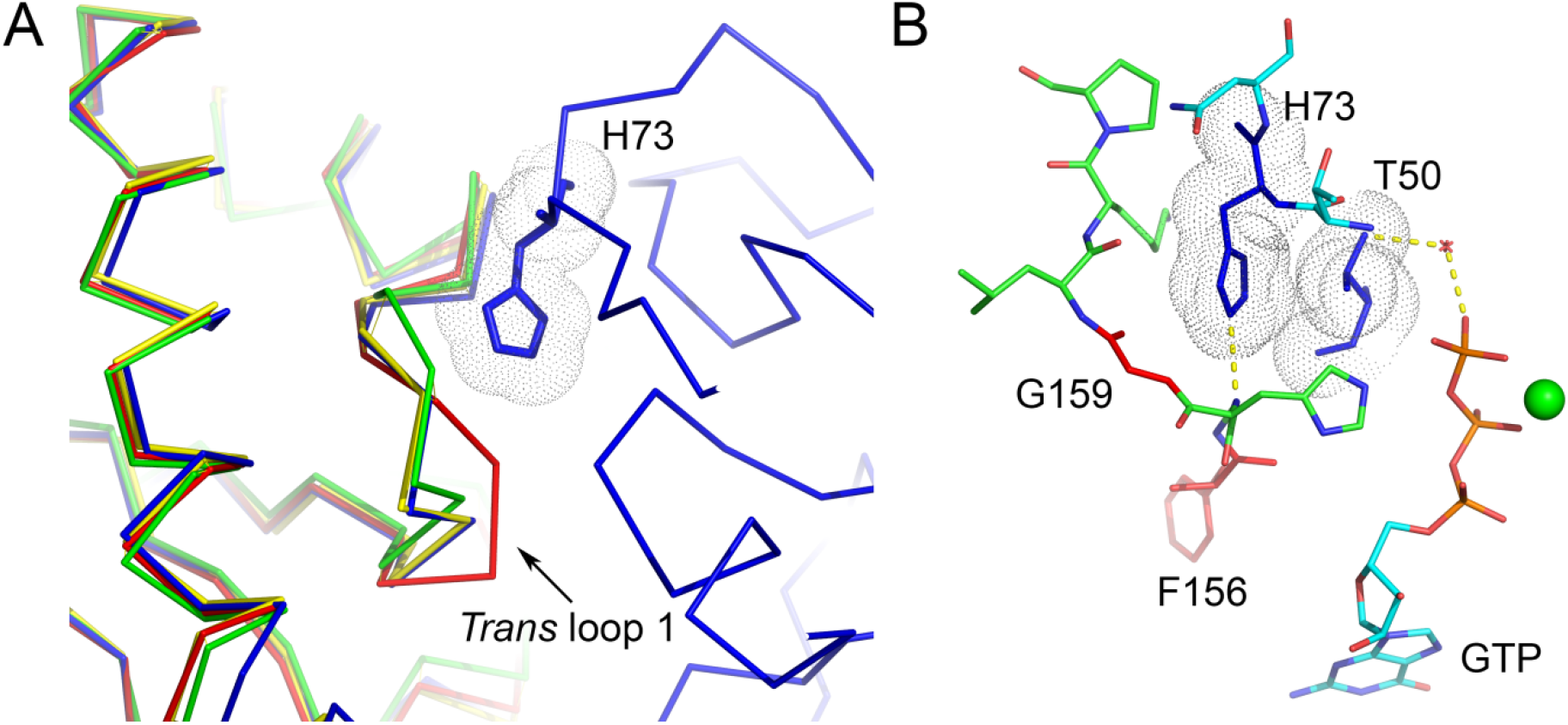
The trans loop 1 and His73. (a) A different conformation is adopted by SEPT2 *trans* loop 1 (red) compared to other subgroups (following the colour scheme used in Fig. 2). (b) Details of the chain of interactions which leads from the γ phosphate of the GTP bound to SEPT11 to the *trans* loop 1 of SEPT2. This includes a water molecule and two *characteristic* residues found in SEPT6/8/11 (His74/His76/His73 (from switch I) and Thr51/Thr53/Thr50 (from the P-loop)), shown in dark blue. The *trans* loop 1 (green) includes the *characteristic* residue Phe156, together with Gly159 (both shown in red). Gly159 is conserved in all SEPT2 subgroup members but is also present in SEPT7 and for this reason, is not considered truly *characteristic*.

The conformation of SEPT2 *trans* loop 1 appears to be an intrinsic feature as it is very similar in previously reported structures of isolated SEPT2 (PDB IDs 2QNR and 3FTQ). It is, therefore, reasonable to consider it to be a pre-formed “lock” into which the histidine “key” from SEPT6/8/11 can readily fit. The “key”, on the other hand, is far from being pre-formed since the histidine comes from the switch I region which is disordered in the absence of an appropriate binding partner. Clearly, the structural organization of switch I and the consequent assembly of the interaction motif involving the histidine and *trans* loop 1 are related events which depend on the encounter between septin subunits coming from the correct subgroups. In this case, the *characteristic* residues and unique structural features involved strongly suggest that such an interaction could only occur between members of the SEPT2 and SEPT6 subgroups.

### The importance of the lack of catalytic activity

Fig. 7b shows that besides Phe156, Gly159 and His73, a further *characteristic* residue in the SEPT6 subgroup, Thr50 (Thr51/Thr53 in SEPT6/SEPT8) from the P-loop (Fig. S4), is involved in establishing a chain of interactions connecting the G-interface to the nucleotide-binding site. This chain leads directly from the γ-phosphate of GTP via a water molecule to the side chain hydroxyl of Thr50, whose methyl group orientates the imidazole of His73 to interact with *trans* loop 1 of SEPT2.

The lack of hydrolytic activity displayed by some septin subgroups has long been something of a quandary. In humans, only the SEPT6 subgroup is devoid of catalytic activity. This is due to the lack of a critical threonine residue in the switch I region [27,28]. Consequently, SEPT6 subgroup members are always found bound to GTP. As a result, the γ-phosphate which remains in the binding site should probably be considered a characteristic feature of the SEPT6 subgroup which, along with the *characteristic* residues themselves, is partly responsible for imbuing selectivity into the interface thereby participating in guaranteeing correct filament assembly.

The side chain of Thr50 is maintained appropriately orientated by Asp106 from the same subunit. Other septin subgroups have a serine in place of threonine and the lack of the side chain methyl means that serine is unable to form the hydrophobic contact which orientates His73 towards SEPT2. In other septin structures, including SEPT2 in the complexes reported here and in the homodimer of SEPT7 at high resolution, this serine shows a variety of different conformations, very different to the unique rotamer observed for Thr50 in all three heterodimeric complexes. In summary, we consider the cluster of residues depicted in Fig. 7b to be a specificity hotspot important for determining the correct pairing of septins during filament assembly.

### Switch I of SEPT2 is also critical

On the opposite side of the interface lies switch I of SEPT2, where the *characteristic* residue Ala71 plays a pivotal role. In all three structures, its methyl side chain is inserted into a pocket formed by Leu161, Ser163, Leu166 and Glu203 of SEPT11 (Leu162, Ser164, Leu167 and Glu204 in SEPT6) as shown in Fig. 8. In other subgroups, this alanine is either lost altogether due to a five-residue deletion in the SEPT6 subgroup or substituted by a residue with a larger side chain. In the former case, the contact would no longer exist and in the latter, the cavity would be too small to accommodate the side chain.

**Fig. 8.**
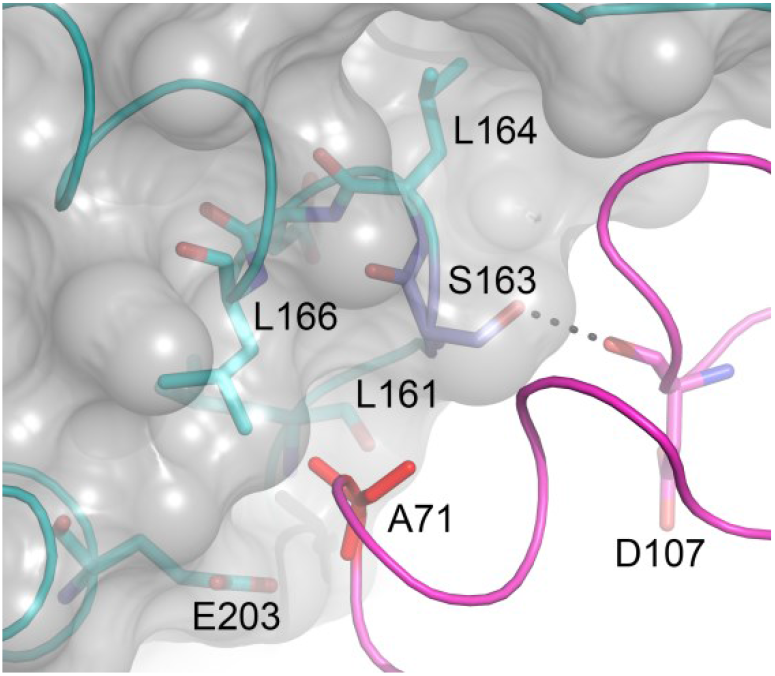
Switch I of SEPT2. The hole-like pocket formed by residues Leu161, Ser163, Leu166 and Glu203 (SEPT11) into which the sidechain of Ala71 from SEPT2 nestles. Ser163 forms a H-bond across the interface with Asp107 in SEPT2. Ala71 (bright red) is a *characteristic* residue of the SEPT2 subgroup and Ser163 (dark blue) can be considered such for the SEPT6 subgroup (only SEPT10 has a Thr at this position). These residues are in Van der Waals contact across the interface. The *characteristic* residues cluster together at the interface to form a specificity hotspot.

Ser163 which lines the pocket and with which Ala71 forms Van der Waals contacts, can be considered *characteristic* of the SEPT6 subgroup although it is conservatively substituted by Thr in SEPT10 (Fig. S3). The hydroxyl group of the Ser163 side chain forms a hydrogen bond with the main chain of Asp107 in SEPT2 (presumably Thr in SEPT10 could do the same). In all other human septins, this residue is a proline which would be unable to form the H-bond. In the structure of SmSEP10 (PDB ID 4KVA), the best available to model the SEPT6 subgroup in the absence of its correct binding partner, the serine points out of the interface. Thus, it would seem that ordering of the SEPT2 switch I region forces Ser163 to reorientate in order to avoid steric hindrance and, in so doing the H-bond to Asp107 of the neighbour becomes feasible. The result is to generate a complementary fit for the *characteristic* Ala71 which is further stabilized by a hydrogen bond formed between its main chain amide and the side chain of Glu203.

In summary, as was observed for the case of the SEPT6/8/11 switch I region, characteristic residues cluster together at the interface to form a specificity hotspot which could only exist with the correct pairing of septin subgroups.

### Switch II in the SEPT2-SEPT6/8/11 complexes

On average, the two switch II regions contribute approximately 300 Å^2^ to the total buried surface area of the G-interface in the SEPT2-SEPT6/8/11 complexes and 380 Å^2^ in the case of SEPT7-SEPT3_T282Y_. In all of the heterodimers described here, the switches form an anti-parallel wide-type β-bridge across the G-interface. This has been described previously [24,25], and it has been recently suggested that this is a characteristic feature of a physiological G-interface and that it distinguishes these from promiscuous ones [20]. This hypothesis is entirely borne out by the four structures reported here. It is also consistent with the fact that none of the heterodimers present β-strand slippage [28,29], a phenomenon which we propose to be incompatible with the formation of the β-bridge and therefore only observed at non-physiological interfaces [20].

Before and after the β-bridge, the switch II region forms an impressive sequence of classical β-turns stabilized by internal main chain hydrogen bonds. Many of these are type II turns which include a Gly, Asp or Asn residue at the *i+2* position [34,35]. The β-bridge, which has been fully described in the homodimer of SEPT7, is part of this sequence (Fig. 9a). It gains additional stability due to Asn123 which uses its side chain to form hydrogen bonds to the main chain of Ala120. However, in the case of the complexes made by SEPT2, the situation is rather different because the homologous position to Asn123 is occupied either by Cys111 in SEPT2 or by a Lys in members of the SEPT6 subgroup. Indeed this lysine is a *characteristic* residue of the SEPT6 subgroup and is surprisingly buried on complex formation. Of the 85 Å^2^ of surface area which Lys111 in SEPT6 (Lys113/Lys110 in SEPT8/SEPT11) would expose if it were monomeric, 60 Å^2^ are buried on forming the dimer with SEPT2. It is one of the most buried lysines in the entire complex which, together with the fact that it is characteristic of the subgroup, is highly suggestive of an important role, be it structural or functional (Fig. 9b). It is of interest to note that recently a novel covalent bond formed between cysteine and lysine side chains has been reported in a series of crystal structures [36,37] but the electron density in the structures reported here provides no direct evidence for such a bond. On the other hand, the complexes with SEPT6 and SEPT8, show clear evidence for the formation of a disulphide bridge between Cys111 and Cys114, albeit of partial occupancy. Further studies of this region of the molecule are clearly required for a fuller understanding of the significance of these observations.

**Fig. 9.**
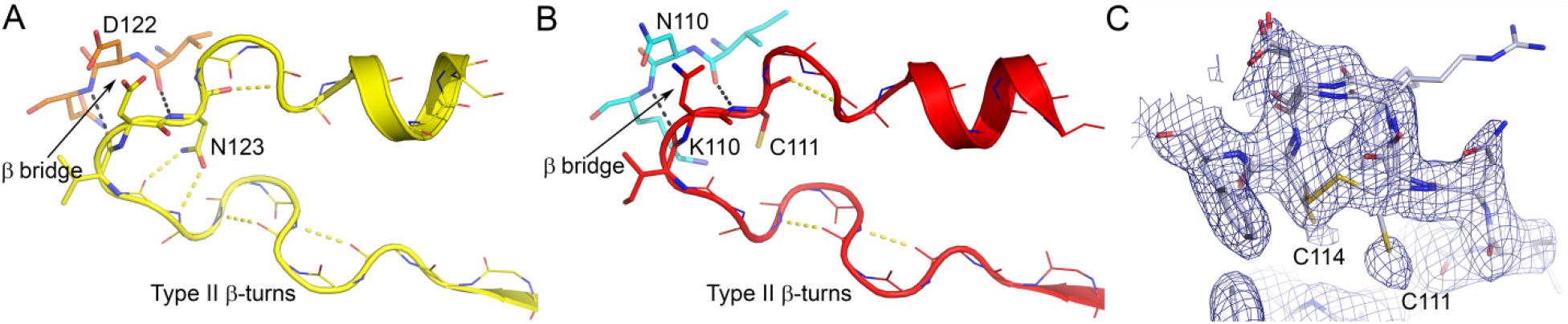
Switch II region. (a) Switch II of SEPT7 at the homodimeric G-interface (PDB ID 6N0B). The two subunits are shown in different shades of yellow (b) Switch II of SEPT2 (red) and SEPT11 (blue) at the heterodimeric G-interface. The residues involved in the wide-type β-bridge across the interface are shown as sticks and the two H-bonds which define the bridge are shown as black dashes. All remaining side chains have been removed for clarity. Hydrogen bonds which stabilize the β-turns are also indicated. In SEPT7 additional hydrogen bonds are formed between the side chain of Asn123 and the main chain. In SEPT2 this residue is replaced by a Cys and in SEPT11 by a Lys. These two side chains are buried beneath the β-bridge. (c) Electron density map (2Fo-FC at 1σ) for the SS Bond formed at switch II.

### Switch II in the SEPT7-SEPT3_T282Y_ complex

It has been reported previously that Phe129 of the switch II region assumes a number of different conformations in the many SEPT3 subgroup structures which have been solved to date [23]. In the heterodimer presented here, where SEPT3 is paired with its definitive G-interface partner (SEPT7), it adopts yet another orientation. Fig. S8a shows the variety of different rotamers for Phe129 seen in the 18 independent subunits reported by Castro et al. On the other hand, Fig. S8b shows that the position adopted in the heterodimer corresponds to none of these but rather is identical to that seen previously for structures which possess a physiological G-interface (SEPT2 and SEPT6/8/11 from the heterodimers reported here and the homodimeric structure for SEPT7). This is related to the formation of the full structure of switch II in terms of its β-turns (Fig. 9) as can be seen from the regularity of the main chain in Fig. S8b in contrast to Fig. S8a. This would suggest that the conformation of the aromatic side chain is dependent on the necessary complementarity at the G-interface with the consequent ordering of switch II and the formation of the inter-subunit β-bridge. It has been speculated that Phe129 may play a role in transmitting information from the G- to the NC-interface on nucleotide hydrolysis [23]. Indeed, the conformational flexibility of this residue, which becomes fixed once the physiological G-interface forms, suggests that not only its conservation in septin sequences but also its dynamics may be of functional significance, in accordance with this proposal.

### The homodimeric NC-interface of SEPT3 is observed in the closed conformation

As mentioned above the NC-interface formed in the SEPT2-SEPT6/8/11 complexes is non-physiological and will not be considered further here. In the case of the SEPT7-SEPT3_T282Y_ structure, the two crystallographically independent NC-interfaces are very different. The 7-7 NC-interface is in the canonical “open” conformation and is stabilized by the typical salt-bridges described previously [23,25]. Since copies of SEPT7 are not expected to interact via an NC-interface in the context of a canonical hexameric or octameric core particle it is probably fair to treat this interface as promiscuous (Fig. 3).

The NC-interface formed between two copies of SEPT3 is physiological and resides at the centre of the octameric particle (Fig. 3a). Previously reported structures from the SEPT3 subgroup have shown a diversity of relative positions for two subunits across this interface, implying an intrinsic malleability not observed for the remaining subgroups. The SEPT3-SEPT3 NC-interface in the SEPT7-SEPT3_T282Y_ complex is observed to be in the closed conformation [23]. This means that the α6 helices are closer together and the α2 helices further apart compared with when the interface is open. The PB2 (polybasic 2) region [38] wraps around α6 of its neighbour, as seen in previous structures of SEPT3 alone, and reaches down deeper into the interface compared with SEPT7 (Fig. 10a). Three new salt bridges arise as a consequence: Arg162 with Glu240, Lys164 with Asp247 and Arg165 with Glu243 (Fig. 10b). These involve residues of the polyacidic region involving helix α5’ and the preceding loop and, as seen in all previous structures of the SEPT3 subgroup, this helix is less inclined with respect to the filament axis than in SEPT7 (Fig. 10a). Together with the projection of PB2 deeper into the interface, this favours the formation of the three salt bridges described above. That formed between Lys164 and Asp247 is particularly interesting as it involves Asp247, a *characteristic* residue in the SEPT3 subgroup. Although its partner, Lys164, is not strictly *characteristic* as defined, only the SEPT3 subgroup has a positively charged residue two positions after the conserved Arg162. This places Lys164 in a unique position to reach down towards the polyacidic region of the neighbouring subunit to interact with Asp247 in a way which other subgroups would be unable. This interaction is therefore specific for an NC-interface formed between two copies of a SEPT3 subgroup member, being unable to be present with any other combination. We expect that the unique positions of PB2 and α5’, resulting in the salt bridges involving *characteristic* residues, would contribute significantly to ensuring that this interface is favoured during spontaneous filament assembly.

**Fig. 10.**
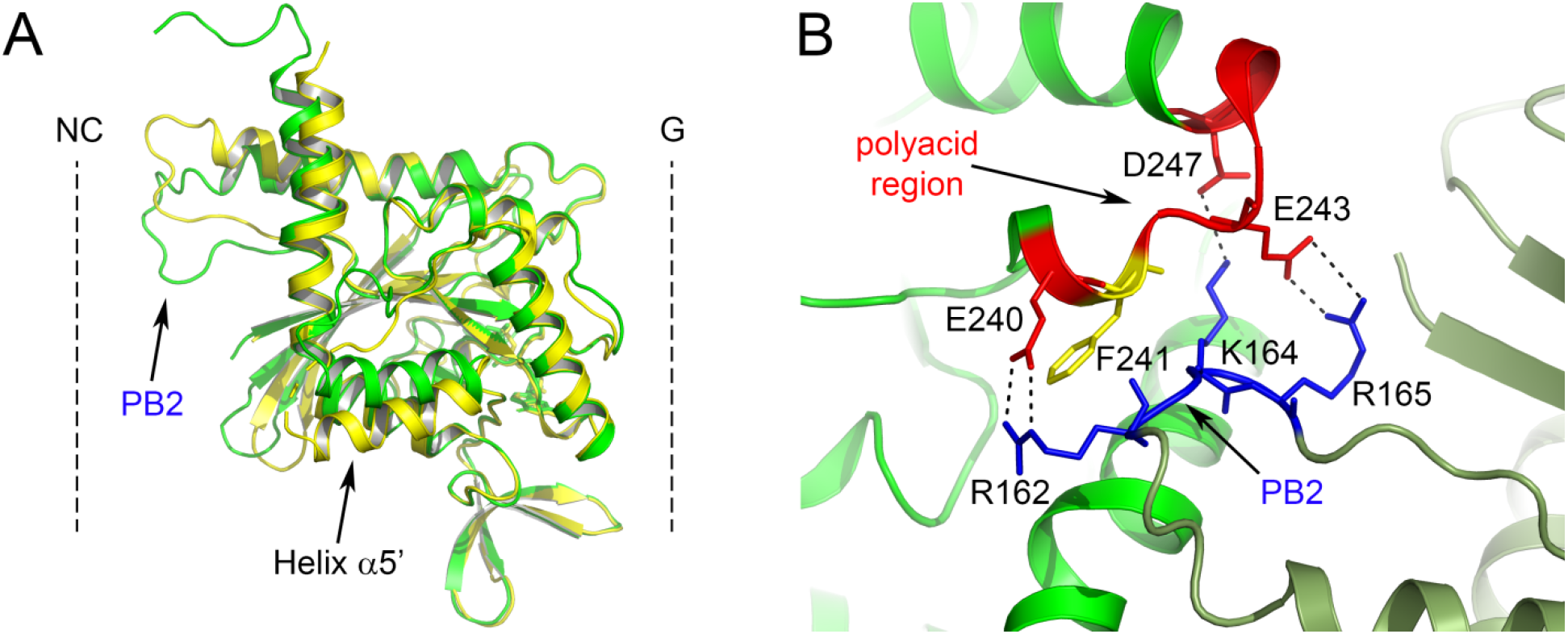
The NC-interface in SEPT7-SEPT3_T282Y_. (a) On comparing SEPT3 (green) with SEPT7 (yellow) from the heterodimeric complex, it can be seen that the PB2 region of the former drops lower into the NC-interface in order to interact with the polyacidic region of its neighbour. (b) Shows the specific salt bridges formed between PB2 (blue) and the polyacidic region (red) of two different SEPT3 subunits (shown in different shades of green) across the closed NC-interface. Asp247 is *characteristic* of the SEPT3 group and Lys164 can be considered so (see text). Phe241 (yellow) is also *characteristic* and aids in holding the polyacidic region and the α5’ helix in an orientation which permits the formation of the salt bridges.

## Conclusion

In the current work, we have made considerable progress towards understanding how the specificity, necessary for correct filament assembly, arises at the two different G-interfaces within the octameric core particle. We have provided the first atomic details of the heterotypic interactions made by the G-domains in both SEPT2-SEPT6 and SEPT7-SEPT3. By substituting SEPT6 by either SEPT8 or SEPT11 it is possible to observe almost complete conservation of the interface contacts providing a structural explanation for Kinoshita’s postulate of the substitutability of septins within a subgroup.

Correct assembly of the oligomer requires that SEPT2 must interact specifically via its G-interface with a SEPT6 subgroup member. This selectivity appears to be guided by the existence of specific interactions between these septins involving unique features of the two subunits involved. Two selectivity hotspots have been identified which include the participation of *characteristic* residues and structural features on either side of the interface, thus guaranteeing that only members of the SEPT2 and SEPT6 subgroups would be able to generate such contacts. The formation of these hotspots requires the ordering of the switch I region in both subunits. These are therefore key players in heterodimer formation as they are responsible for a significant part of the interfacial buried surface and imbue the complex with a thermostability which is less evident in that formed between SEPT7 and SEPT3, where the switch regions barely participate in inter-subunit contacts.

With the atomic detail now available for all of the G-interfaces present within both the octameric (SEPT2-SEPT6 and SEPT7-SEPT3) and hexameric (SEPT2-SEPT6 and SEPT7-SEPT7) core particles, some consistent themes begin to emerge. Most notable is that they all have completely ordered switch II regions, part of which forms an antiparallel wide-type β-bridge. This now appears to be well established as a signature of a physiological G-interface. It is associated with the formation of a series of β-turns (which complete the switch II region) and also with fixing a conserved aromatic residue (Phe129 in SEPT3) into a single and well-defined conformation. Although we have speculated previously as to the potential relevance of this in communication between adjacent interfaces, its veracity and mechanism remain to be established.

Slowly, a picture of the details of specific septin interfaces is emerging. Once those which have so far resisted the efforts of structural biologists have also been fully characterized, the spontaneous assembly of septin hexamers, octamers and filaments should become more readily understandable.

## MATERIALS AND METHODS

### Cloning, expression, and purification

The G domains of SEPT2 (SEPT2G) and SEPT6 (SEPT6G) were produced as described previously in Kumagai et al., 2019 with minor modifications. The co-expression of members from the SEPT2 subgroup (SEPT2G, SEPT4G and SEPT5G) and SEPT6 subgroup (SEPT6G, SEPT8G and SEPT11G) were performed by inserting the CDS for the corresponding G domains into the bicistronic expression vector pETDuet-1 (Novagen). The G domain of SEPT7 (SEPT7G) was expressed as described in Brognara et al., 2019. Finally, the constructs to co-express SEPT7G with SEPT9 (SEPT9GC) or SEPT3 (SEPT3GC and SEPT3GC_T282Y_) were based on those constructs used in Castro et al., 2020 and Macedo et al., 2012, respectively, but here using the pETDuet-1 vector, to allow for pairwise co-expression. The details of all constructs used are described in Table S1.

*Escherichia coli* BL21 Rosetta™(DE3) (Merck/Novagen) cells harbouring the co-expression vectors were grown at 37 °C in LB medium (Lysogeny-Broth) supplemented with ampicillin (50 μg·ml^−1^), chloramphenicol (34 μg·ml^−1^), and when applicable kanamycin (30 μg·ml^−1^). At a cell density corresponding to an absorbance of 0.6-0.8 at 600 nm, the culture was cooled to 18 °C and the protein co-expression induced by 0.3 mM isopropyl 1-thio-β-D-galactopyranoside. After 16 h, cells were harvested by centrifugation at 10,000 g for 40 min. at 4 °C, and suspended in lysis buffer (Table S1). After cells lysis by sonication, the soluble fraction was isolated by centrifugation at 16,000 g for 1h at 4 °C and then loaded onto a 5 mL HisTrap HP column (GE Healthcare) previously equilibrated in the lysis buffer. Subsequently, the column was washed with 10 vol. of IMAC elution buffer (Table S1) and the proteins were eluted using a linear gradient from 10-100% of imidazole in the same buffer. A Superdex 200 10/300 GL column (GE Healthcare) pre-equilibrated in SEC buffer (Table S1) was used to perform the size exclusion chromatography (SEC) as the final purification step. The desired concentration of the heterodimers for each experiment was then achieved by performing cycles of concentration at 800 g, using an Amicon Ultra centrifugal filter (Merck/Millipore) with a 30 kDa cutoff.

### Size exclusion chromatography coupled with multi-angle light scattering (SEC-MALS)

The oligomeric state of the complexes was evaluated using a miniDAWN TREOS three-angle light scattering detector (Wyatt Technology) and Optilab T-rEX refractometer (Wyatt Technology). This system was coupled to a Waters 600 HPLC system (Waters) for the size exclusion chromatography. A volume of 50 μl of each sample at 3 mg·ml^−1^ was loaded onto a Superdex 200 Increase 10/300 GL column (GE Healthcare) equilibrated in 25 mM Hepes pH 7.8, 300 mM NaCl and 5 mM MgCl_2_. The data collection and analysis were performed via ASTRA 7 software (Wyatt Technology).

### Nucleotide identification

GTP or GDP present in the purified complexes was detected by the method described by [39] with minor modifications. Samples at a concentration of 25 μM were incubated with ice-cold HClO_4_ (final concentration 0.5 μM) for 10 mins. Samples were then centrifuged at 20,000 g for 10 min. at 4 °C to separate the denatured protein pellet. The supernatant was transferred to a new microtube and neutralized with ice-cooled solutions of KOH 3 M (1:6), K_2_HPO_4_ 1 M (1:6) and 0.5M acetic acid. Samples were stored at −20 °C for at least 1 hour, then thawed and centrifuged (20,000 g for 10 min. at 4 °C) for HPLC analysis. The nucleotides were separated by anion exchange chromatography on a Protein Pack DEAE-5 PW 7.5 mm × 7.5 cm column (Waters), pre-equilibrated in 25 mM Tris-HCl pH 8.0, coupled to an Alliance 2695 chromatography system. Samples of 200 μL were loaded and eluted with a linear NaCl gradient (0.1 – 0.45 M in 10 min.). The column was calibrated with separate 200 μL samples of GDP and GTP at 10 μM in the column buffer. Absorbance was monitored at 253 nm.

### Thermal shift assay

The thermal stability of the dimers was assessed using intrinsic fluorescence measurements with a Tycho NT.6 instrument, NanoTemper Technologies [40]. The temperature was ramped within the range from 35 to 90 °C at a rate of 30 °C/min and the intrinsic fluorescence ratio (330/350 nm) of each sample at 5 μM in 25 mM Hepes 7.8, 300 mM NaCl, 5mM MgCl_2_ and 5% (v/v) glycerol, was monitored and used to calculate the midpoint unfolding inflection temperature (Ti). Standard 10 μl capillaries were used and the data were measured in triplicate and analysed using the equipment software.

### Crystallization, data collection and structure determination

Dimeric complexes were crystallized by the sitting drop vapour diffusion method using the Morpheus screening kit (Molecular Dimensions). A drop of 0.2 μl of fresh purified protein was mixed with 0.2 μl of the reservoir solution consisting of: 100 mM Bicine/Trizma base pH 8.5, 12.5% (w/v) PEG 1000, 12.5% (w/v) PEG 3350, 12% (w/v) MPD and 20 mM of each of the following, D-glucose, D-manose, D-galactose, L-fucose, D-xylose and N-acetyl D-glucosamine (for the SEPT2G-SEPT6G complex at 2 mg·ml^−1^); 100 mM MES/imidazole pH 6.5, 10% (w/v) PEG 8000, 20% (w/v) ethylene glycol and 20 mM of each of the following, 1,6-hezanediol, 1-butanol, (RS)-1,2,-propanediol, 2-propanol, 1,4-butanediol and 1,3-propanediol (for the SEPT2G-SEPT8G at 4 mg·ml^−1^); 100 mM MES/imidazole pH 6.5, 10% (w/v) PEG 20000, 20% (w/v) PEG 550 MME and 30 mM of each of the following, sodium nitrate, disodium hydrogen phosphate and ammonium phosphate (for the SEPT2G-SEPT11G at 4 mg·ml^−1^); 100 mM MES/imidazole pH 6.5, 10% (w/v) PEG 4000, 20% (v/v) glycerol and 20 mM of the following, sodium formate, ammonium acetate, sodium citrate tribasic dihydrate, potassium sodium tartrate tetrahydrate and sodium oxamate (for the SEPT7G-SEPT3GC_T282Y_ complex at 23 mg·ml^−1^). After 24h at 291 K, the crystals were harvested and cryo-cooled in liquid nitrogen for data collection.

X-ray diffraction data were collected at 100 K on the Diamond Light Source using beamline I24 housing a PILATUS3 6M detector (SEPT2G-SEPT6G, SEPT2G-SEPT11G and SEPT7G-SEPT3GC_T282Y_) or beamline I04 with an Eiger2 × 16M detector (SEPT2G-SEPT8G). The data were indexed, integrated and scaled using xia2 [41] and the structures were solved by molecular replacement with Phaser [42]. To solve the SEPT2-SEPT6 complex the individual monomers of human SEPT2 (PDB ID 2QNR) and *Schistosoma* Septin 10 (PDB ID 4KVA) [28] were used. To solve the SEPT2-SEPT11 complex, the refined structure of SEPT2-SEPT6 was used as the search model, whereas for SEPT2-SEPT8, the structure of SEPT2-SEPT11 was employed. Finally, SEPT7-SEPT3_T282Y_ was solved using individual monomers of SEPT3 (PDB ID 4Z54) [23] and SEPT7 (PDB ID 6N0B) [20]. Alternate rounds of refinement and model rebuilding using Phenix [43] and Coot [44] yielded the final models. The data collection, refinement statistics and PDB code are summarised in Table S3. Contact residues across the G-interface were determined with DIMPLOT [45,46] and subject to minor manual adjustments after visual inspection. The surface areas of the interacting regions and their specific amino acids were calculated using PISA [47]. Figures were generated with Pymol v 2.05.

## ACESSION NUMBERS

Coordinates and structure factors have been deposited in the Protein Data Bank with accession numbers **<u>6UPA</u>, <u>6UPQ</u>, <u>6UPR</u> and <u>6UQQ</u>**.

The sequences and respective coordinates for each of the proteins is readily available on NCBI and UniProt repository under the codes as provided by Table S1.

## ACKNOWLEDGEMENTS

We gratefully acknowledge the support of the São Paulo Research Foundation (FAPESP) via grants 2014/15546-1 and 2016/04658-9, along with CNPq via PIBIC grants 144121/2016-6 and 164288/2017-1, and also CAPES via PROEX program. We are also grateful to Susana Sculaccio, Andressa Alves Pinto and Derminda Isabel de Moraes for excellent technical support. We recognize the essential role played by DIAMOND–UK in providing access to PX-beamlines and also to their technical staff during synchrotron data collection. We are extremely grateful to Daniel Maturana and Carolina Carriel from NanoTemper Technologies for providing access to the Tycho instrument for the thermal stability measurements.

## AUTHOR CONTRIBUTIONS

H.V.D.R., D.A.L., G.B. and H.M.P performed the experiments. H.M.P. and J.B. coordinated and undertook the X-ray data collection and processing. R.C.G., A.P.U.A. and H.M.P conceived the study and supervised its execution. All authors participated in analysing the data and in writing and revising the manuscript.

## DECLARATION ON INTEREST

The authors declare no competing interests.

## SUPPLEMENTARY MATERIAL SUPPLEMENTARY MATERIAL

### Supplementary figures

**Fig. S1.**
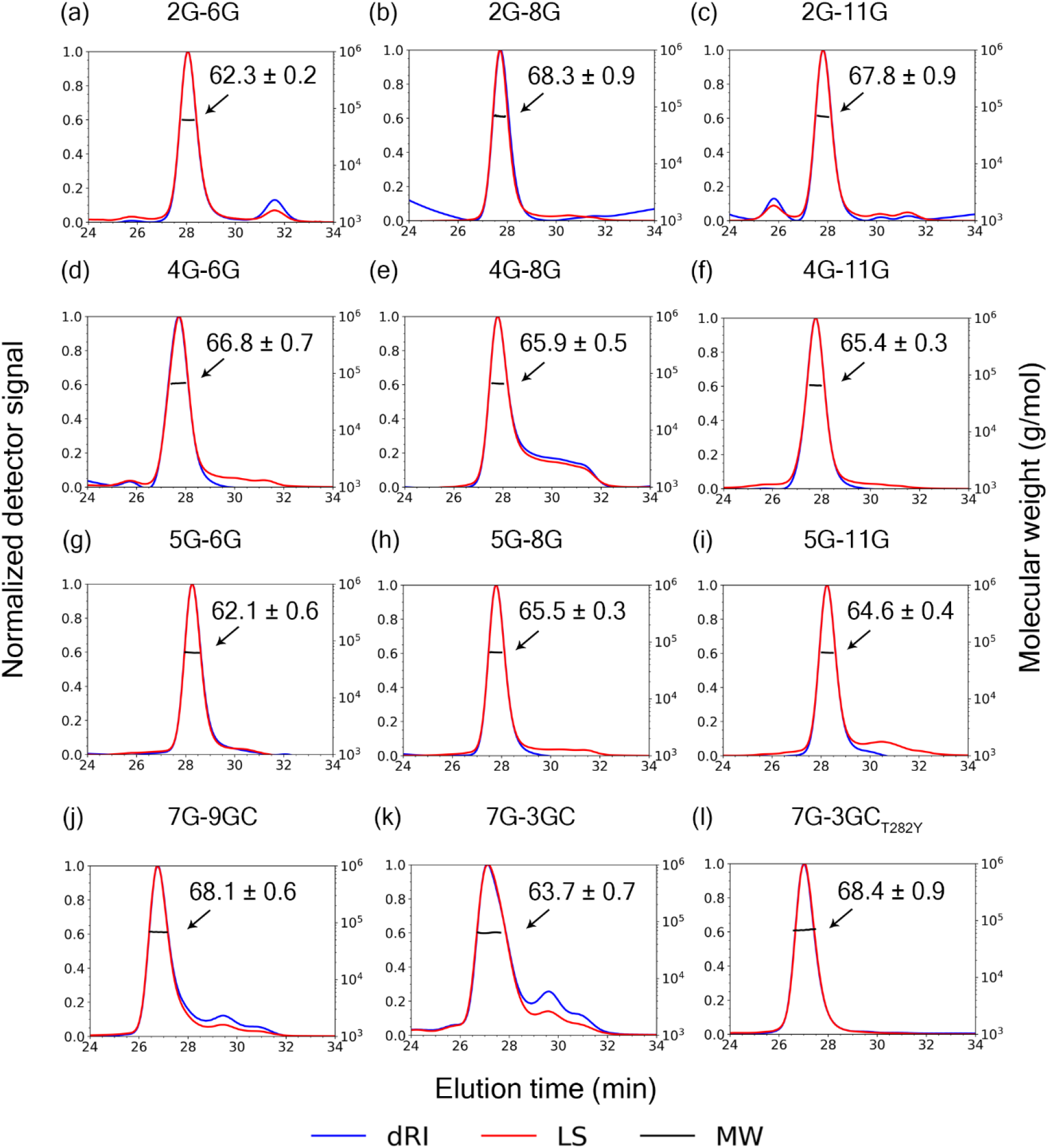
SEC-MALS traces for all of the heterodimeric complexes. In all cases curves corresponding to the change in the normalized differential refractive index (blue), normalized scattered light intensity at 90°(red) and calculated molecular weight of the corresponding peak (black) are given.

**Fig. S2.**
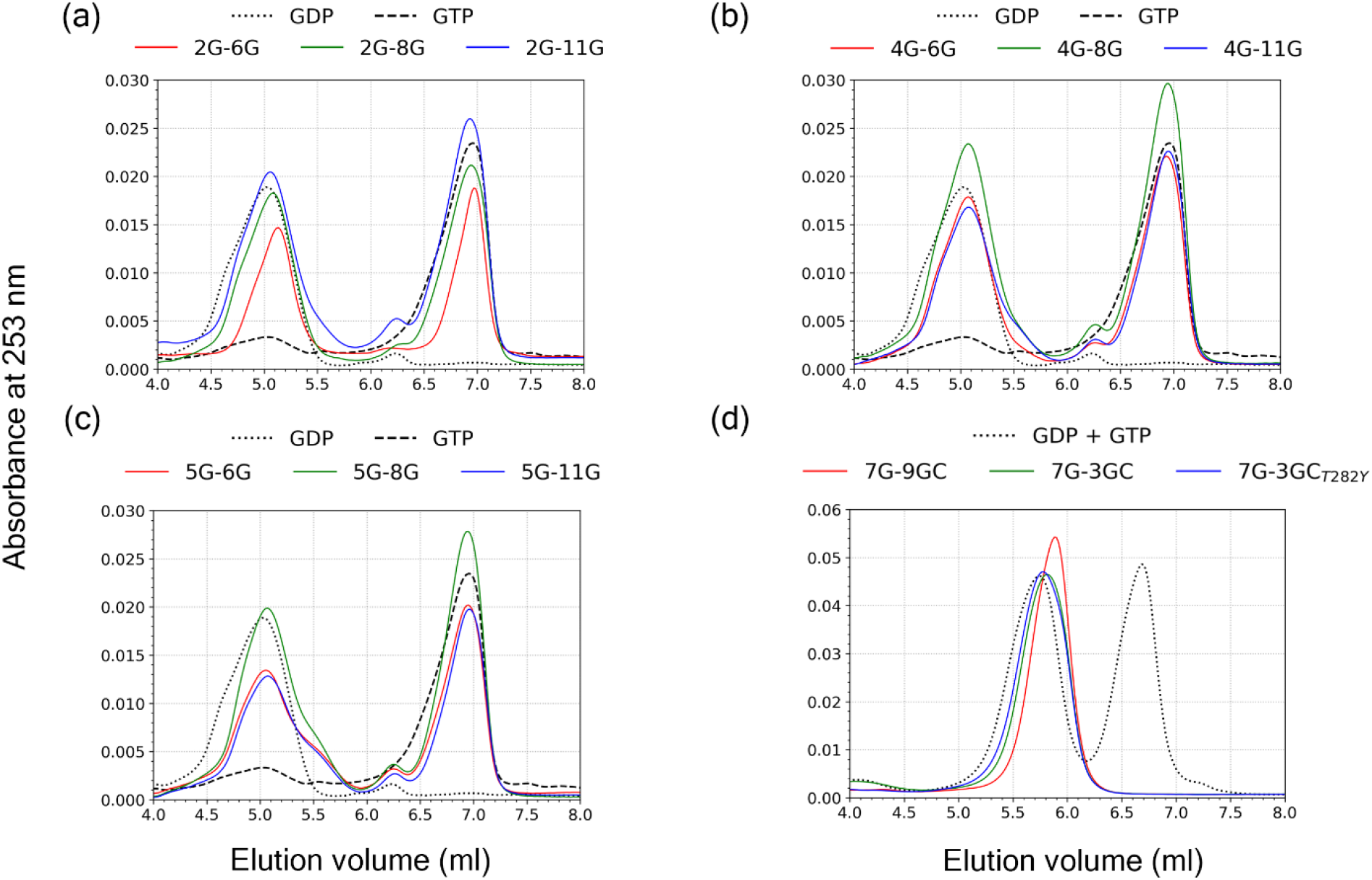
Nucleotide content for the 12 heterodimeric complexes. Each panel shows the analysis performed on three heterodimers together with the traces for GDP and GTP. All dimers show the presence of both nucleotides with the exception of those containing a SEPT3 subgroup member (SEPT3 or SEPT9).

**Fig. S3.**
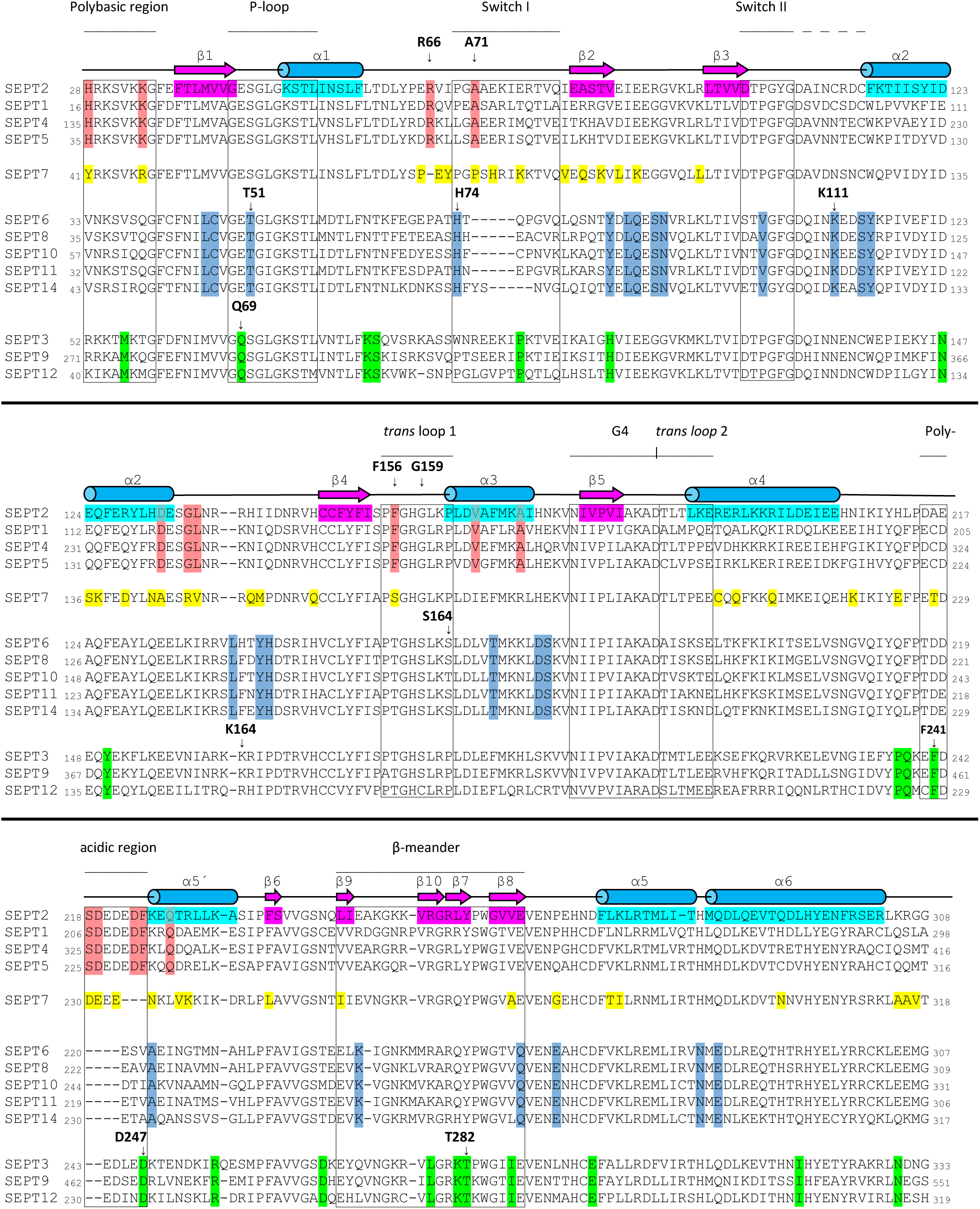
Alignment of the G-domains of all 13 human septins, divided into the four established subgroups. The first sequence in each subgroup is considered to be representative and is responsible for the name (SEPT2, SEPT7, SEPT6 and SEPT3). Elements of secondary structure are indicated by horizontal bars, based on the structure of SEPT2 in the complex with SEPT6 (β-strands are shown as magenta arrows and α-helices as light blue cylinders). Structural features which are relevant to the interfaces, including the switch regions, G motif, *trans* loops, β-meander, polybasic and polyacidic regions are also indicated. Residues highlighted correspond to so-called *characteristic* residues, namely those that are completely conserved within the subgroup but absent from all other sequences. These represent a “fingerprint” of the subgroup and are shown in red (SEPT2) blue (SEPT6) yellow (SEPT7) and green (SEPT3). *Characteristic* residues, which are specifically cited in the text are indicated with an arrow and residue number. For this purpose, the numbers quoted are for the representative sequence of the subgroup. Some additional residues of importance to the discussion which are “almost *characteristic*” are also indicated but without the coloured highlight. The UniProt isoform accession number and initial and final residue coordinates for each of the displayed sequences used to assemble the alignment are as follows: **SEPT2** <u>[Q15019-1]</u>(28-308), **SEPT1**<u>[Q8WYJ6-1]</u>(16-298), **SEPT4**<u>[O43236-1]</u>(135-416), **SEPT5**<u>[Q99719-1]</u>(35-316), **SEPT7**<u>[Q16181-1]</u>(41-318), **SEPT6**<u>[Q14141-1]</u>(33-307), **SEPT8**<u>[Q92599-2]</u>(35-309), **SEPT10**<u>[Q9P0V9-1]</u>(57-331), **SEPT11**<u>[Q9NVA2-1]</u>(32-306), **SEPT14**<u>[Q6ZU15-1]</u>(43-317), **SEPT3**<u>[Q9UH03-2]</u>(52-333), **SEPT9**<u>[Q9UHD8-2]</u>(271-551)**, SEPT12**<u>[Q8IYM1-1]</u>(40-319).

**Fig. S4.**
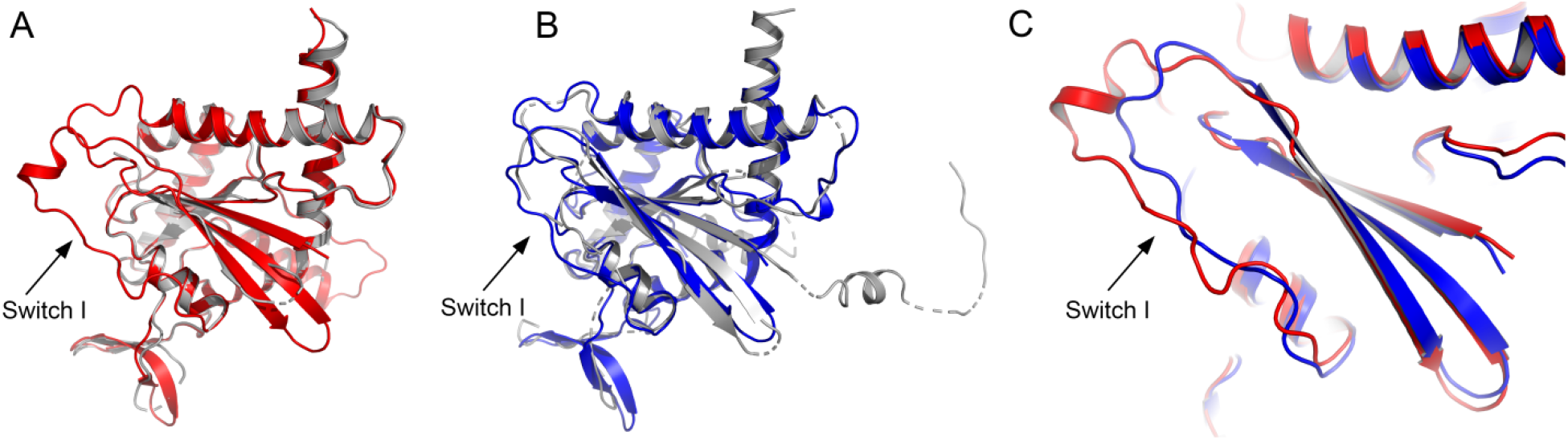
Comparison of individual septins from the best available structures. (a) Overlay of the highest resolution structure for isolated SEPT2 (from PDB ID 2QNR, grey) with SEPT2 from the complex with SEPT6 (red). In both cases, SEPT2 is bound to GDP. (b) The structure of SEPT6-GTP (from the heterotrimer SEPT2-SEPT6-SEPT7, 2QAG, in grey) with SEPT6-GTP as observed in the complex with SEPT2 (blue). (c) Comparison of the switch I region of the two subunits in the SEPT2-SEPT6 complex.

**Fig. S5.**
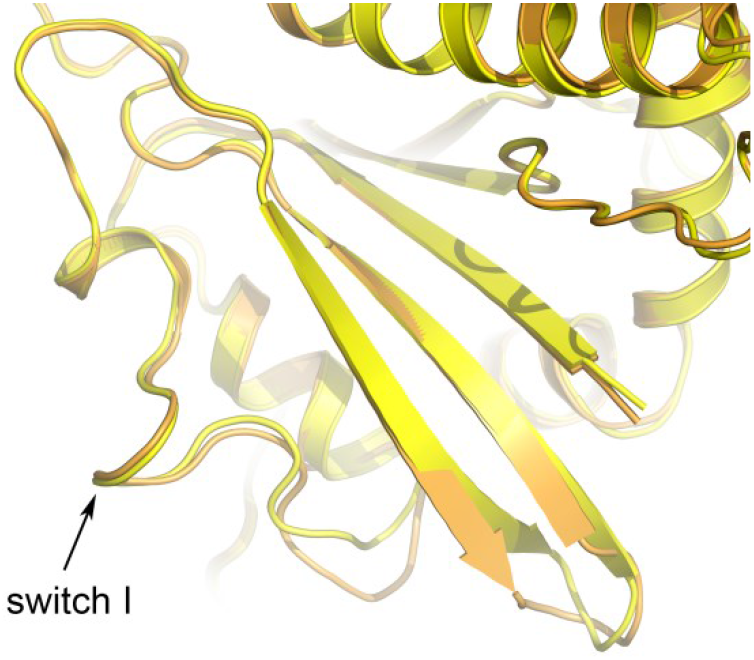
Switch I. Superposition of an ordered switch I region in SEPT7 (6N0B, gold) and in the SEPT7 subunit of the complex with SEPT3_T282Y_(yellow). The conformation of the switch is effectively identical in the two structures.

**Fig. S6.**
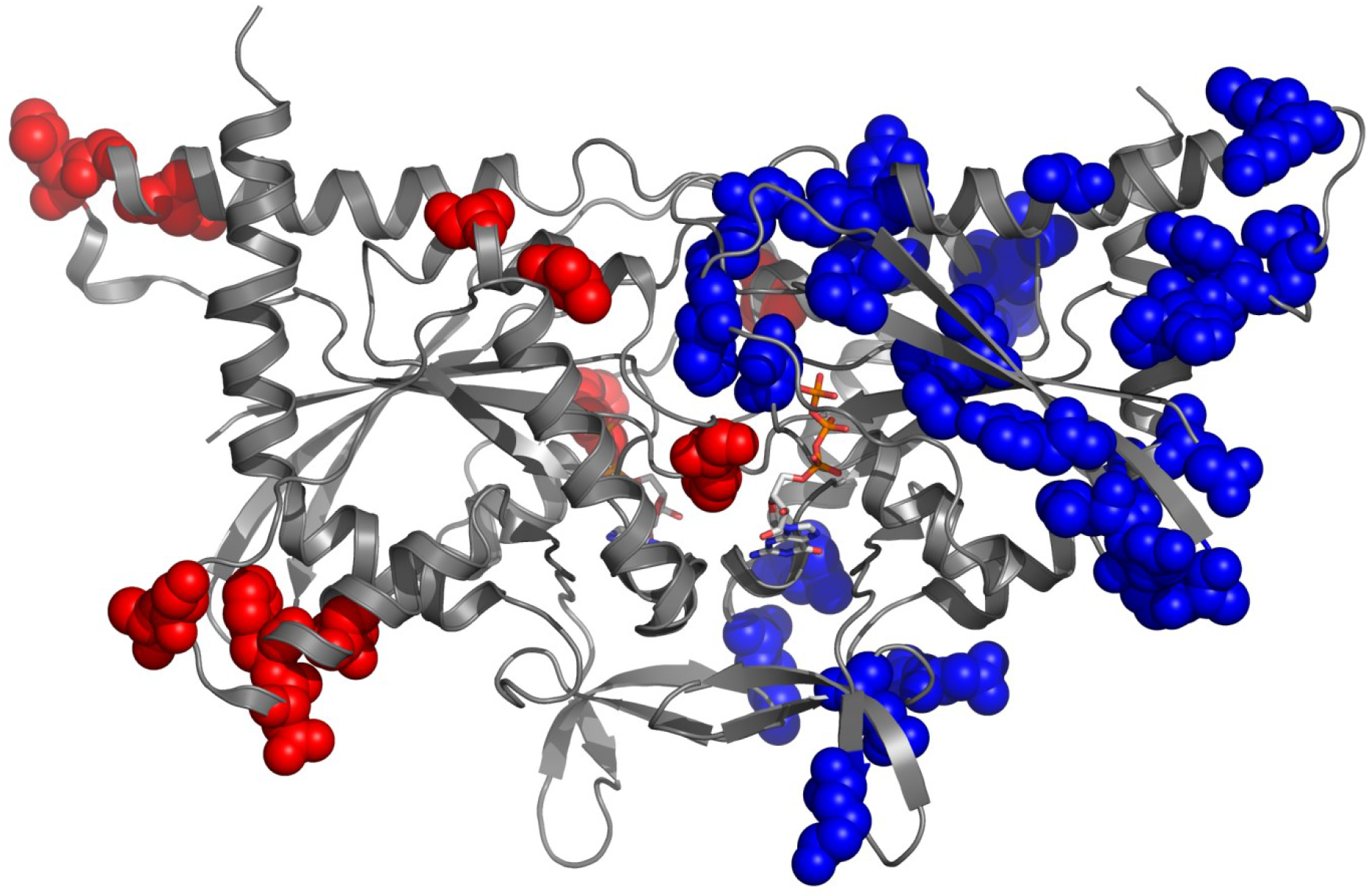
*Characteristic* residues for the SEPT2 and SEPT6 subgroups. The distribution of *characteristic* residues for the SEPT2 and SEPT6 subgroups are shown on the structure of the SEPT2-SEPT11 structure in red and blue respectively. Several of these lie close to the interface.

**Fig. S7.**
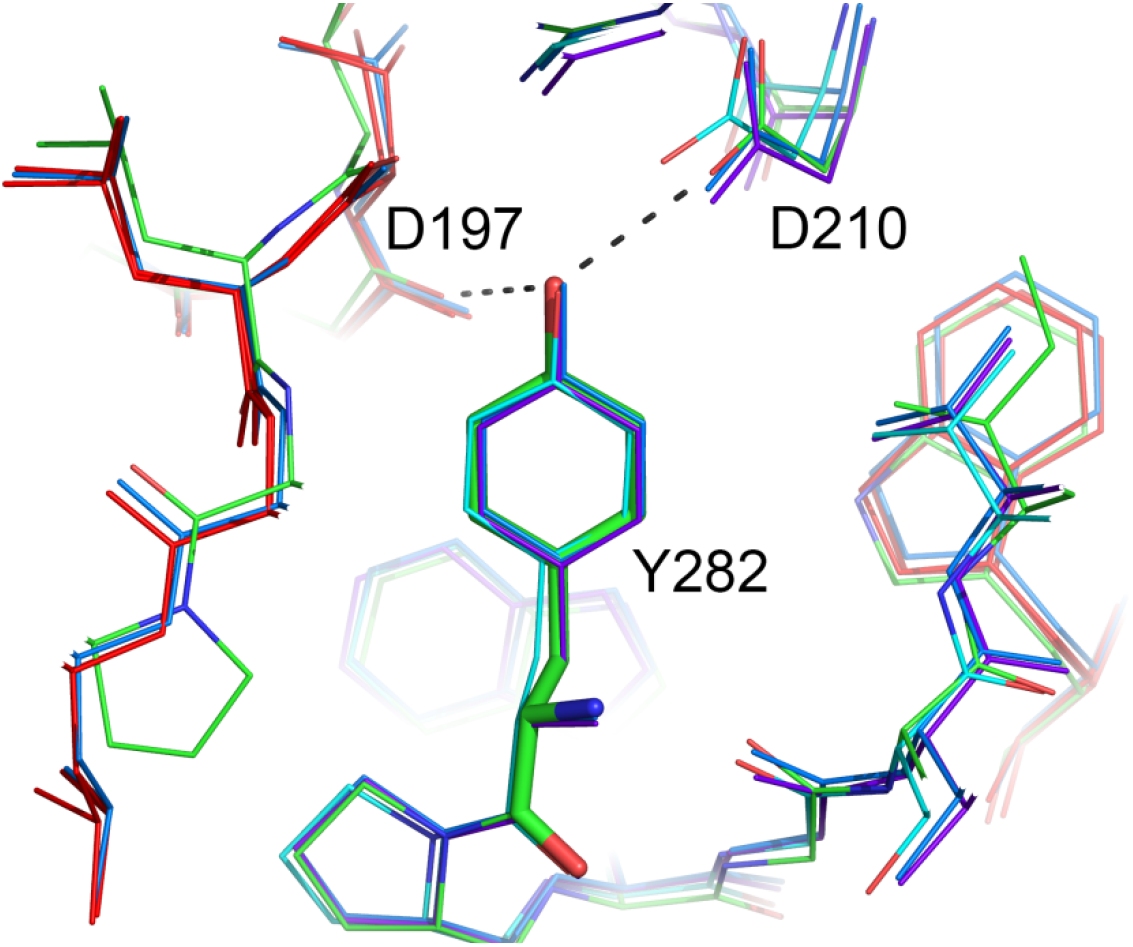
Tyr298 in the SEPT3 mutant. The mutated Tyr282 in the heterodimer SEPT7-SEPT3mut (green) forms the same interactions at the G-interface observed in the heterodimeric complexes formed with SEPT2 including that with Asp197 of SEPT7 (yellow). SEPT6, SEPT8 and SEPT11 are shown in different shades of blue and SEPT2 in red.

**Fig. S8.**
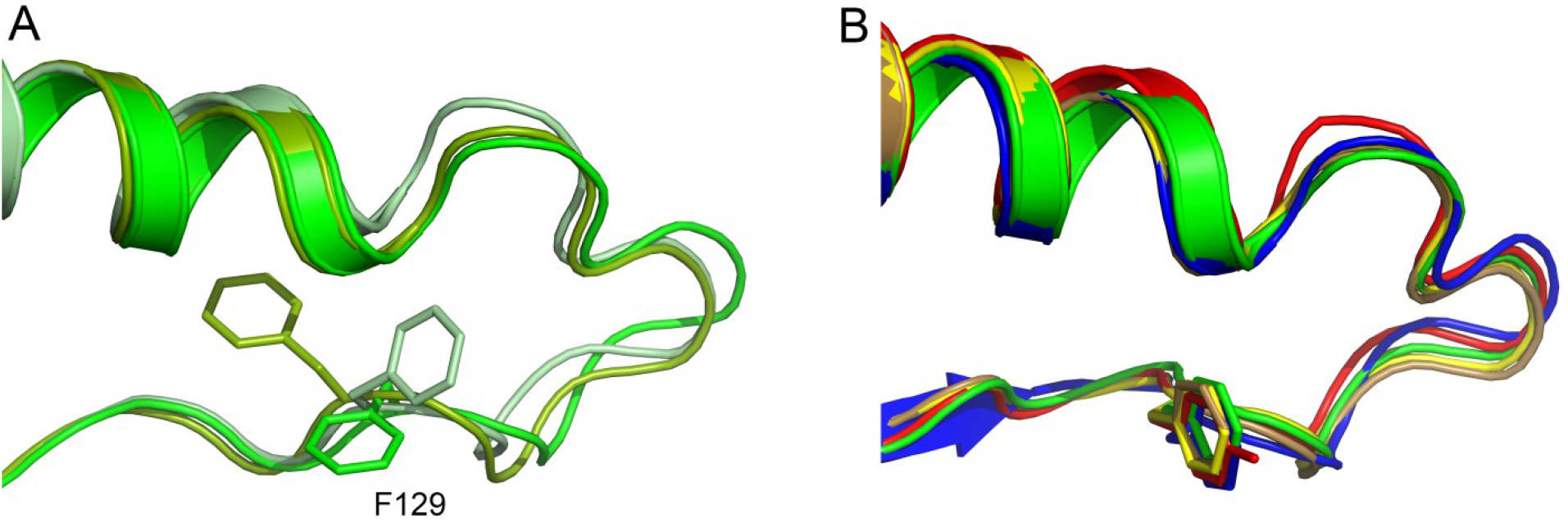
Phe129 of the switch II region of SEPT3. (a) The variety of conformations observed for Phe129 (or its homologue) in different structures of the SEPT3 subgroup when forming a non-canonical homodimeric interface. Three representative examples are shown in different shades of green. (b) In all of the structures in which a canonical interface is formed (including that of SEPT3_T282Y_ in complex with SEPT7), this phenylalanine adopts an identical conformation (different from all those seen in (a)) and the main chain forms the sequence of β-turns (see Fig. 9) SEPT3_T282Y_ (green), SEPT7 (yellow), SEPT2 (red) and SEPT11 (blue) are from the heterodimeric complexes described in this work. SEPT7 (brown) is from the homodimeric structure (PDB ID 6N0B).

## Supplementary tables

**Table S1.**
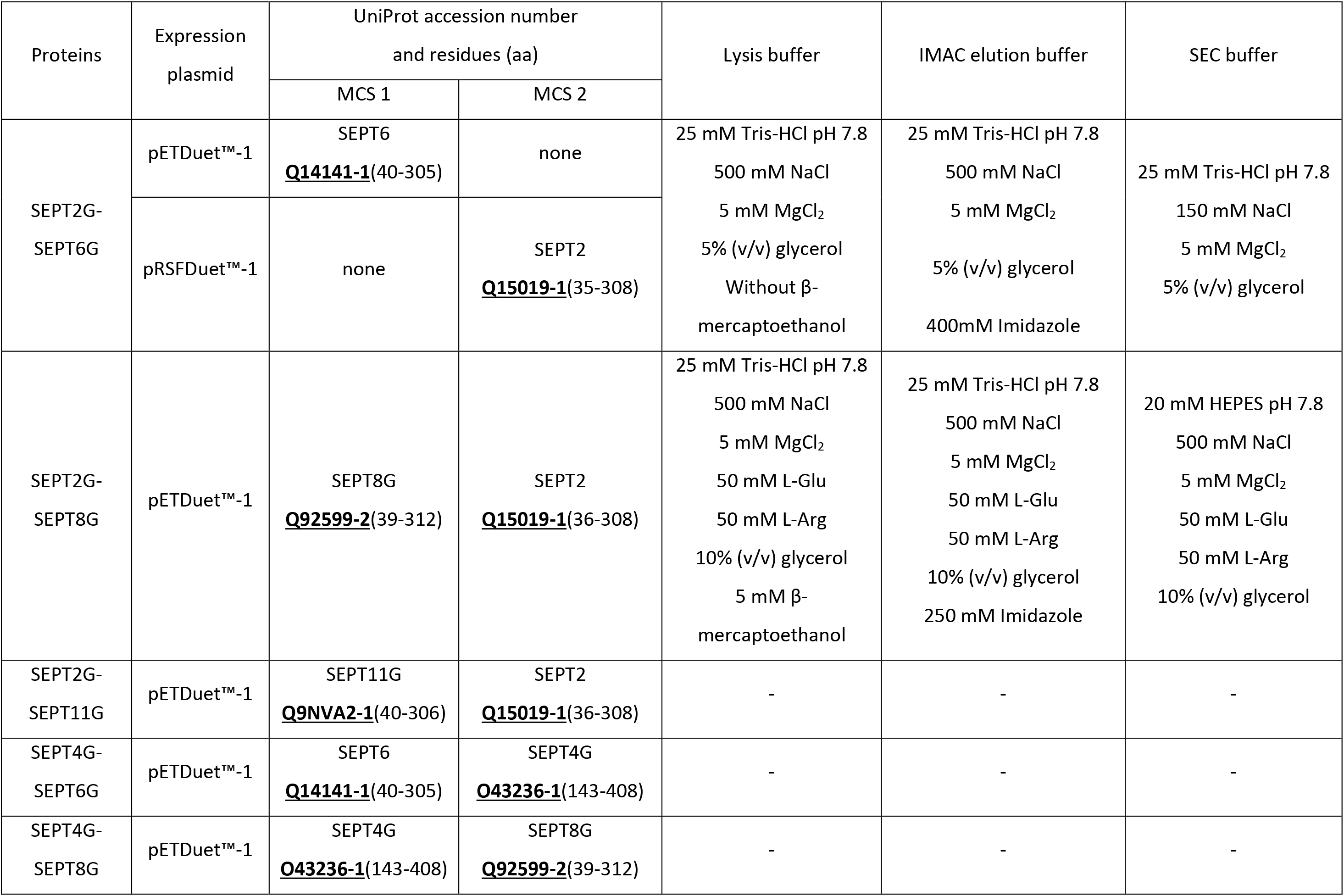
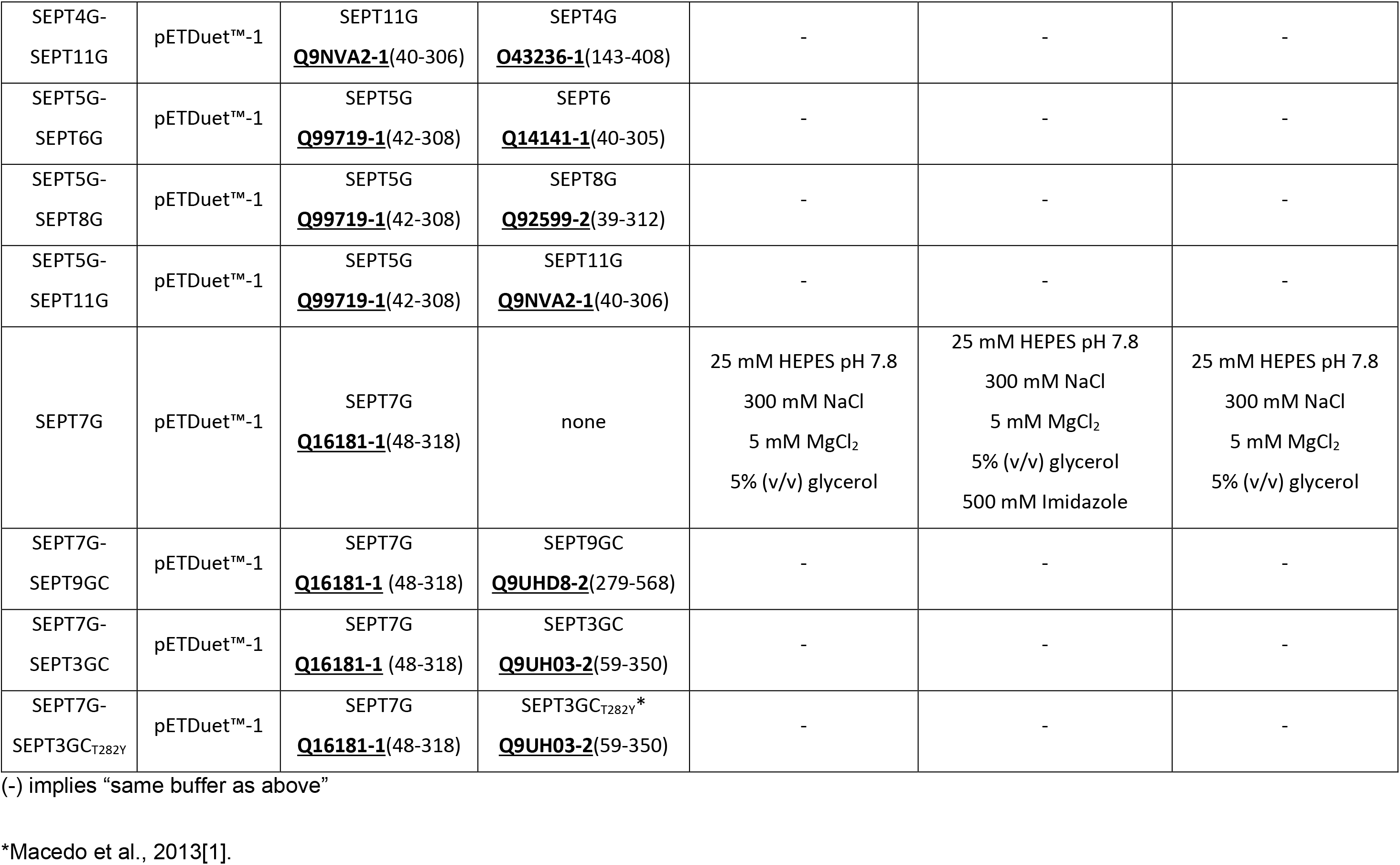
Conditions used for the expression and purification of the dimers described in this work.

**Table S2.** Experimentally determined and theoretical (sequence derived) molecular weights for the heterodimers. The error refers to the deviations in the molecular weights across the peak of the heterodimer and the percentage difference refers to that between the value determined experimentally and that predicted from the amino acid sequence.

| Sample | Molecular weight (kDa) |  | Difference (%) |
| --- | --- | --- | --- |
|  | Sequence predicted | SEC-MALS Determined |  |
| SEPT2G-SEPT6G | 63.7 | 62.3 ± 0.2 | 2.2 |
| SEPT2G-SEPT8G | 63.7 | 68.3 ± 0.9 | 7.2 |
| SEPT2G-SEPT11G | 65.2 | 67.8 ± 0.9 | 4.0 |
| SEPT4G-SEPT6G | 64.3 | 66.8 ± 0.7 | 3.9 |
| SEPT4G-SEPT8G | 64.1 | 65.9 ± 0.5 | 2.8 |
| SEPT4G-SEPT11G | 64.6 | 65.4 ± 0.3 | 1.3 |
| SEPT5G-SEPT6G | 63.7 | 62.1 ± 0.6 | 2.4 |
| SEPT5G-SEPT8G | 65.2 | 65.5 ± 0.3 | 0.4 |
| SEPT5G-SEPT11G | 64.0 | 64.6 ± 0.4 | 0.9 |
| SEPT7G-SEPT9GC | 66.4 | 68.1 ± 0.6 | 2.6 |
| SEPT7G-SEPT3GC | 66.4 | 63.7 ± 0.7 | 4.0 |
| SEPT7G-SEPT3GC<sub>T282Y</sub> | 66.4 | 68.4 ± 0.9 | 3.0 |

## Notes

### Competing Interest Statement

The authors have declared no competing interest.

